## Supplementary material for "Inter-tissue convergence of gene expression during ageing suggests age-related loss of tissue and cellular identity": Figure supplements

**Figure 1-figure supplement 1. Age distribution of samples**

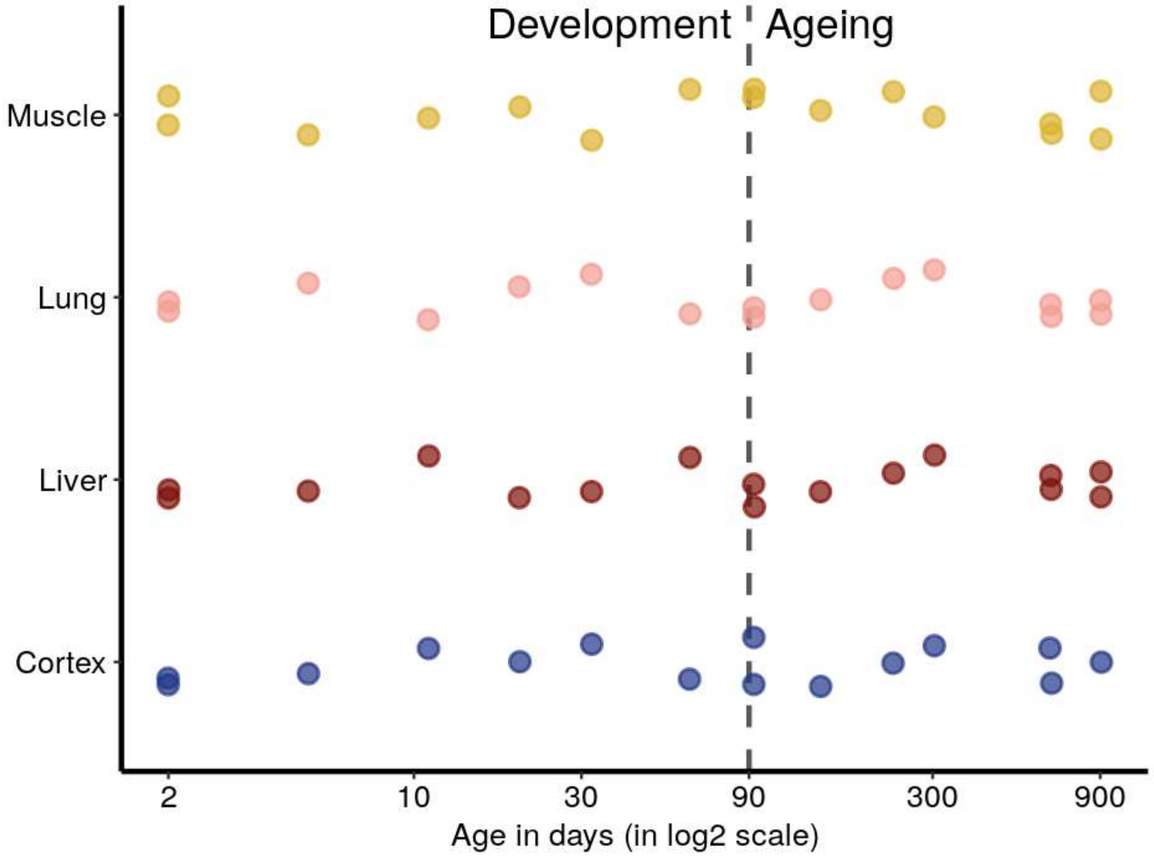


*The x-axis shows the age in days in a log2 scale and the y axis lists different tissues. The period from 2 to 61-days-old mice are considered as postnatal development (referred to as development for brevity in the main text), and above 90-days-old as the ageing period. Random jitter was added on the y-axis to avoid overlap between points.*

**Figure 1-figure supplement 2. PCA with all samples (tissue effect removed)**

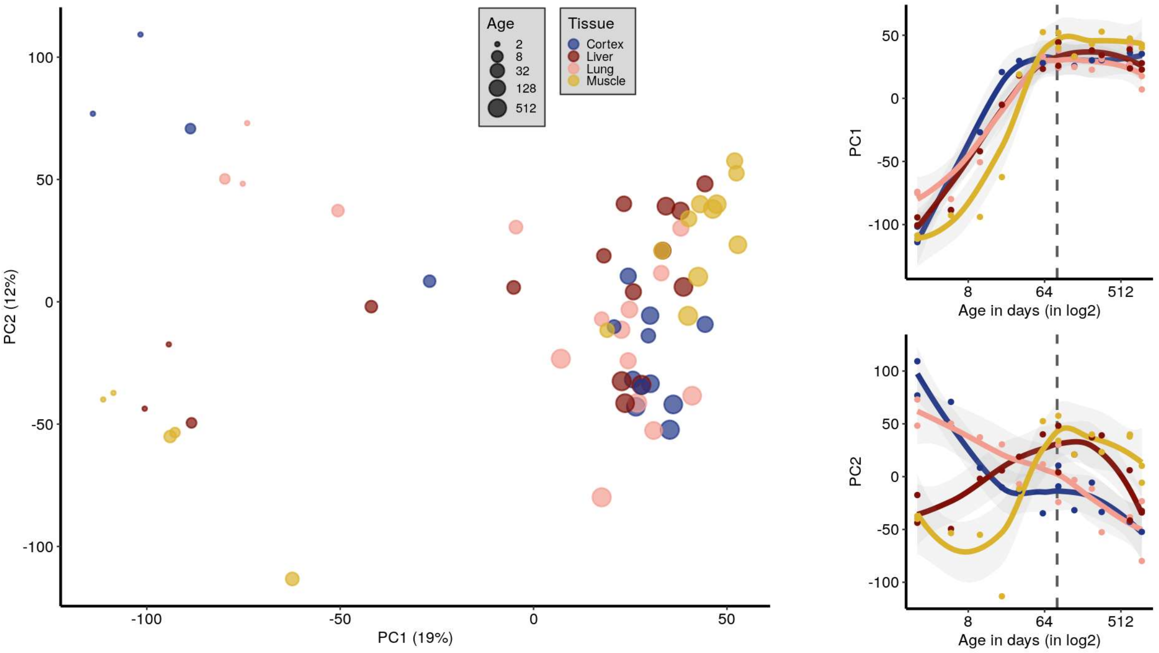


*Principal component analysis (PCA) using all samples (n=16) after each tissue is standardised separately (i.e. gene expression values for individuals are scaled to mean=0, sd=1). PC1 (x-axis) and PC2 (y-axis) are plotted and the variation explained by each PC is denoted within parentheses on each axis. The size of the points indicates the age and the colour shows the tissue. The plots on the right show the correlations between the PCs (y-axis) and age (x-axis, on the log2 scale) in development and ageing. PC1-age Spearman's correlation test during development (n=7 mice); abs($\rho_{dev}$)=[0.88, 0.99], nominal $p_{dev}$<0.01 for each tissue, same test for PC2 vs age; abs($\rho_{dev}$)= [0.30, 0.99], nominal $p_{dev}$<0.01 except muscle (**Figure 1-source data**).*

**Figure 1-figure supplement 3. PCA with development and ageing periods separately**

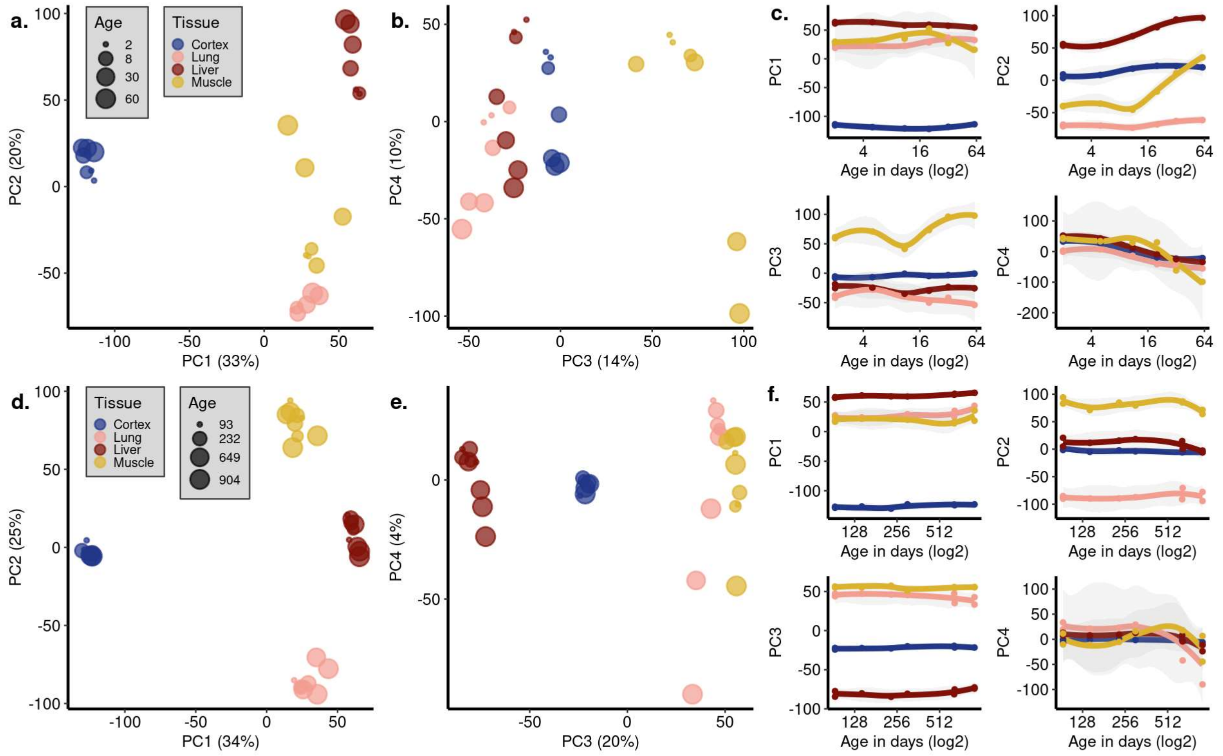


*Principal component analysis (PCA) using only the samples from the development period (2- to 61 days of age, n=7) (**a-c**) and the ageing period (93- to 904 days of age, n=9) (**d-f**). **a,d)** PC1 (x-axis) vs PC2 (y-axis) and **b,e)** PC3 (x-axis) vs PC4 (y-axis) are plotted and the variation explained by each PC is denoted within parentheses on each axis. The size of the points indicates the age and the colour shows the tissue. **c,f)** Correlation between the PCs (y-axis) and age (x-axis, in the log2 scale) in development (c) and ageing (f). **c)** Age-effects can be observed in PC2 and PC4 in development: PC2-age Spearman's correlation test, abs($\rho$)= [0.72, 0.94], nominal p<0.05 in 3/4 tissues; PC4-age Spearman's correlation test, abs($\rho$)= [0.88, 0.99], nominal p<0.01 in all tissues. Inter-tissue transcriptome divergence can be observed as a trend in PC3-PC4 space (change in the mean Euclidean distance among tissues with age in PC1-4 space, $\rho$=0.95, p=0.0008). **f)** A small age-effect can be observed in PC4 in ageing: PC4-age Spearman's correlation test: abs($\rho$)= [0.11, 0.77], nominal p<0.05 in 2/4 tissues. Inter-tissue transcriptome convergence can be observed as a subtle trend in PC1-4 space (change in mean Euclidean distance among tissues with age in PC1-4 spaces, $\rho$=-0.64, p=0.059). All PC-age correlation test results are given in **Figure 1-source data.***

**Figure 1-figure supplement 4. Permutation test results for shared expression trends among tissues**

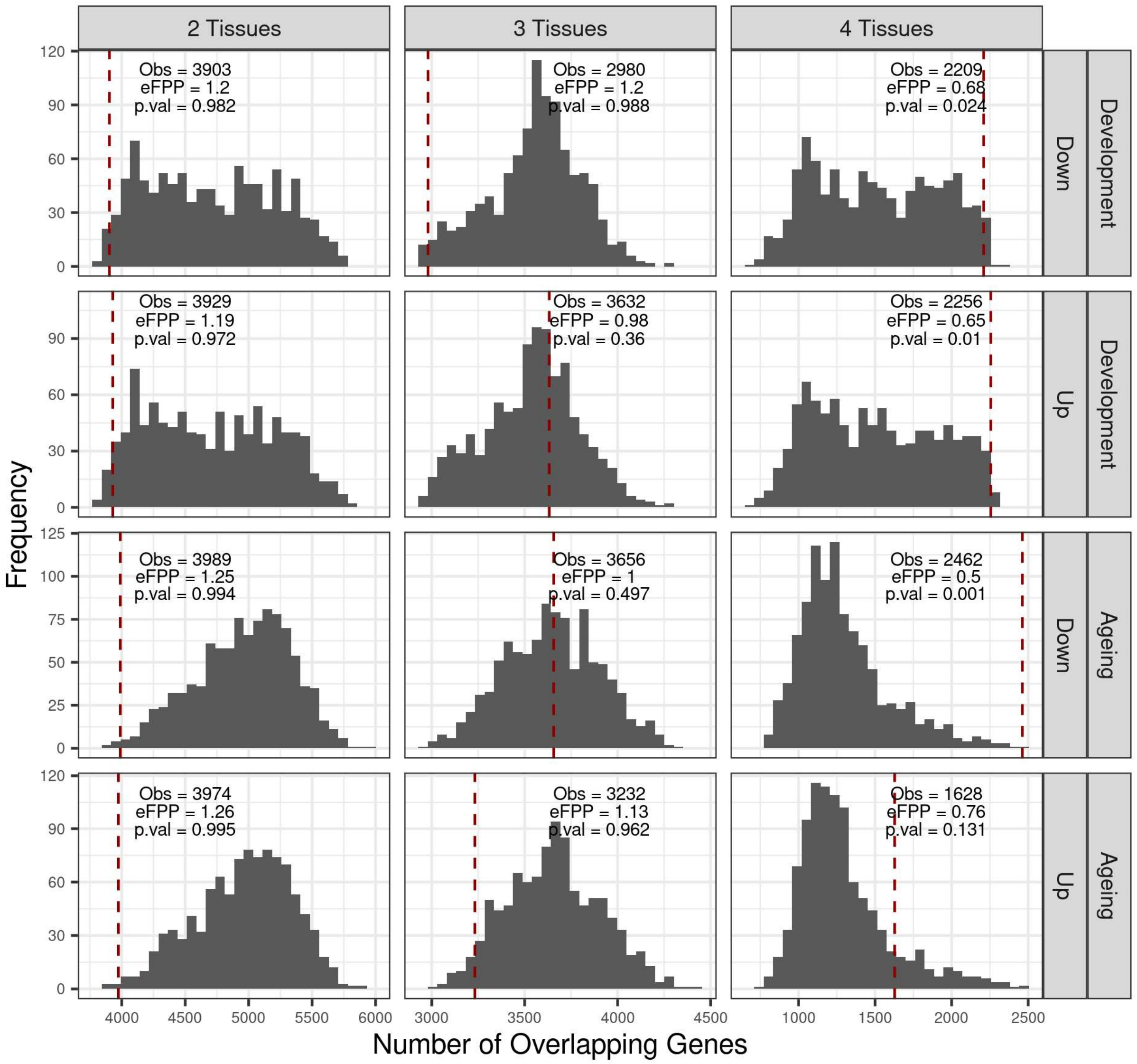


*Permutation test results of shared up/down genes across tissues for development and ageing periods. "Up" and "down" indicate positive and negative expression-age correlations ($\rho$), respectively. No significance cutoff was applied for choosing up/down genes in tissues (i.e. only considering $\rho>0$ or $\rho<0$). The null distributions are created by permuting individual ages and calculating expression-age correlations in each tissue, then summing the number of genes changing in the same direction in 2, 3, and 4 tissues. The red dashed lines show the observed values, also noted as "Obs:". The eFPP (estimated false positive proportion) was calculated as the ratio between the median expected value from the permutations and the observed value. P-values were calculated as the proportion of permutations that are*

*higher than or equal to the observed value.*

**Figure 1-figure supplement 5. Shared age-related genes among tissues in development and ageing**

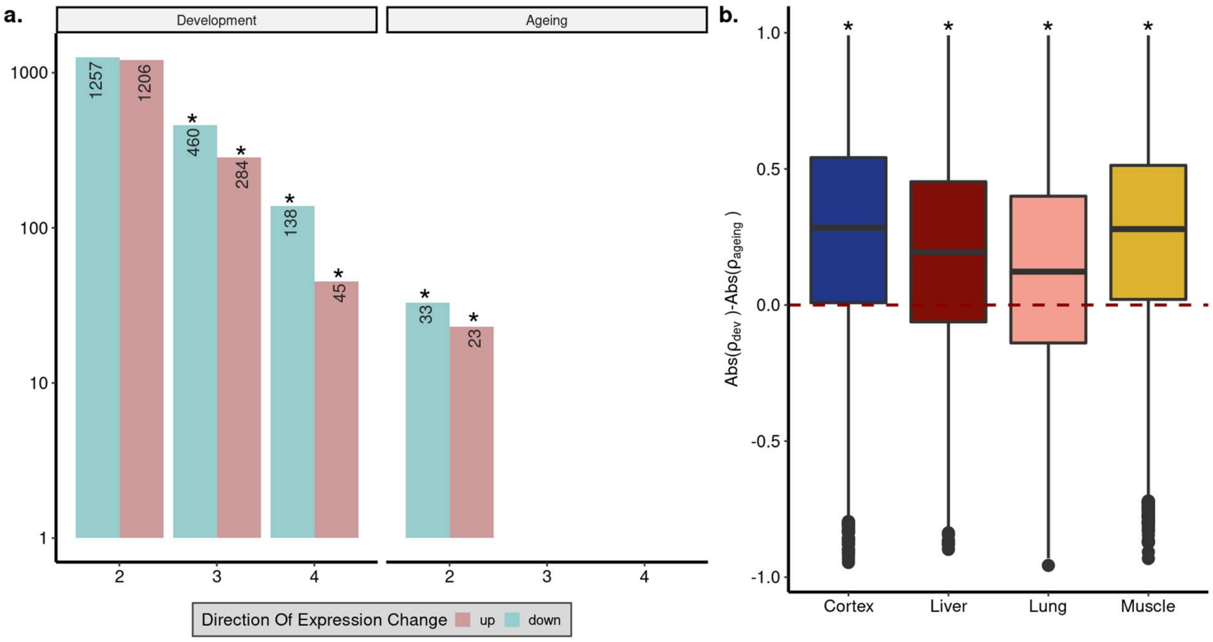


***a)*** *Overlap between significant (FDR corrected p-value<0.1) age-related gene sets among tissues. The x-axis shows the number of tissues compared; 2: overlap in two tissues, 3: overlap in 3 tissues, 4: overlap in 4 tissues. Cyan: down-regulation with age, pink: up-regulation with age. Significant overlaps (permutation test, p<0.05, (see* ***Figure 1–figure supplement 6*** *for test results)) are indicated with asterisks.* ***b)*** *The differences between the magnitude of age-related expression changes in development and ageing: (abs($\rho_{dev}$)-abs($\rho_{ageing}$)), for each gene (n=15,063 genes) in four tissues (Wilcoxon signed-rank test, $p<10^{-16}$ for each tissue).*

**Figure 1-figure supplement 6. Permutation test results for significant trends shared among tissues**

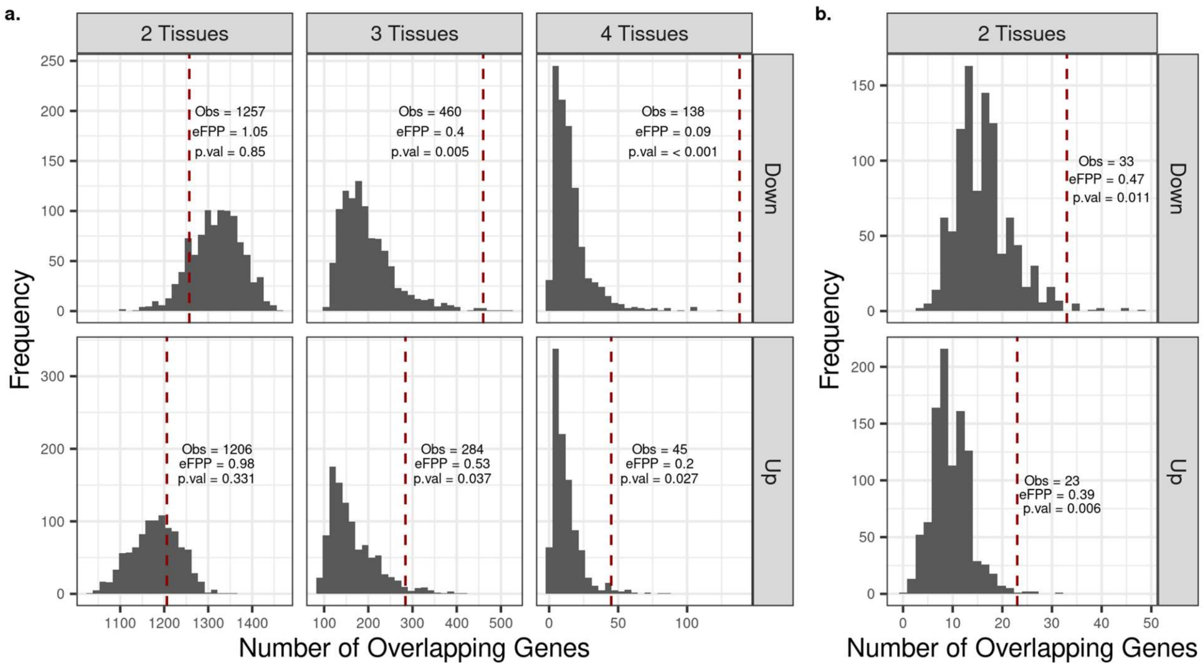


*Permutation test result for shared "up" (or "down") genes among tissues in development* ***(a)*** *and ageing* ***(b)****. "Up" and "down" indicate positive and negative expression-age correlations (ρ), respectively. Significant up/down genes were chosen with FDR corrected p-value<0.1 and their overlap across tissues were calculated. To create the null distributions, we chose as many up (or down) genes in permutations as the observed up (or down) genes in each tissue and then calculated the number of overlapping genes among tissues. The dashed red line shows the observed number of shared up (or down) genes between tissues and eFPP was calculated as the ratio between the median expected value from the permutations and the observed value. "Obs:" number of genes displaying the same significant age-related change pattern among tissues. The p-value was calculated as the proportion of permutations that are higher than or equal to the observed value.*

**Figure 1-figure supplement 7. Similarities between age-related gene expression changes among tissues**

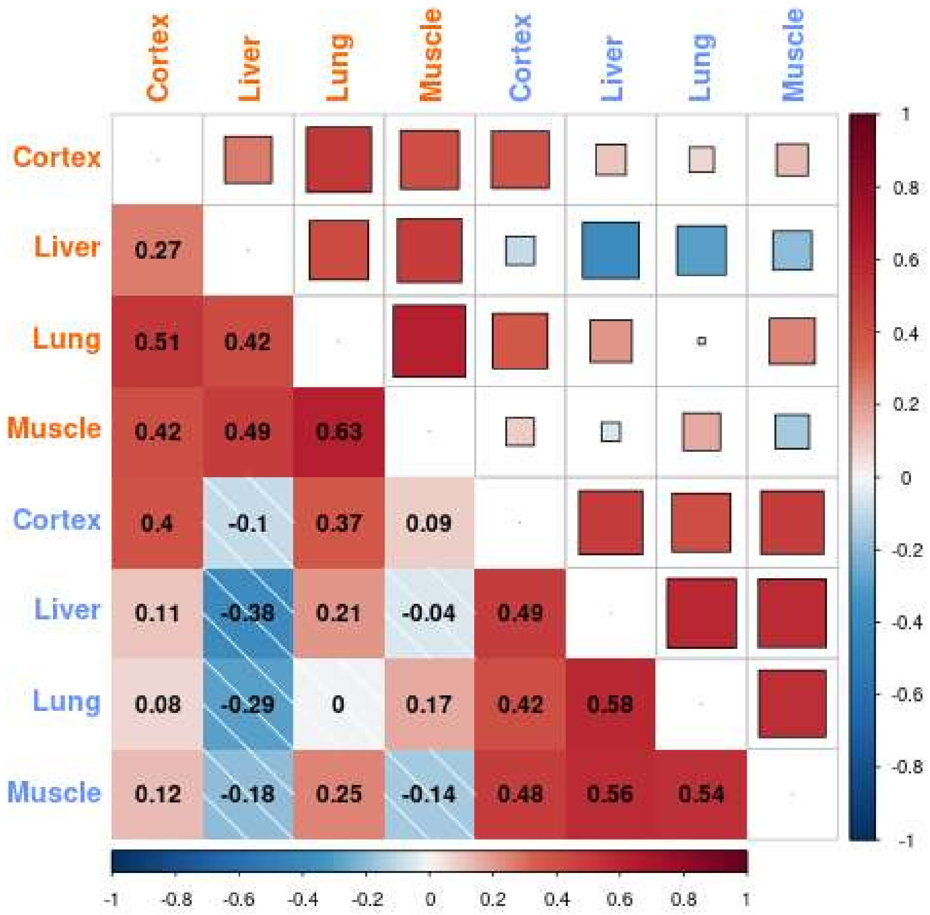


*The similarity between the age-related gene expression changes (Spearman's correlation coefficient between expression and age) across tissues in development and ageing. Similarities were calculated using Spearman's correlations coefficient between expression-age correlations (with cutoff: $|\rho|>0.6$) across tissues. No significance cutoff was used for expression change similarities. The intensity of the colours shows the magnitude of the correlation coefficient, where darker blue indicates a stronger negative correlation and darker red indicates a stronger positive correlation. Correlation values are written on the lower triangle. The colour of the tissue label indicates development (orange) and ageing (blue) datasets.*

**Figure 1-figure supplement 8. Permutation test results for reversal patterns in each tissue**

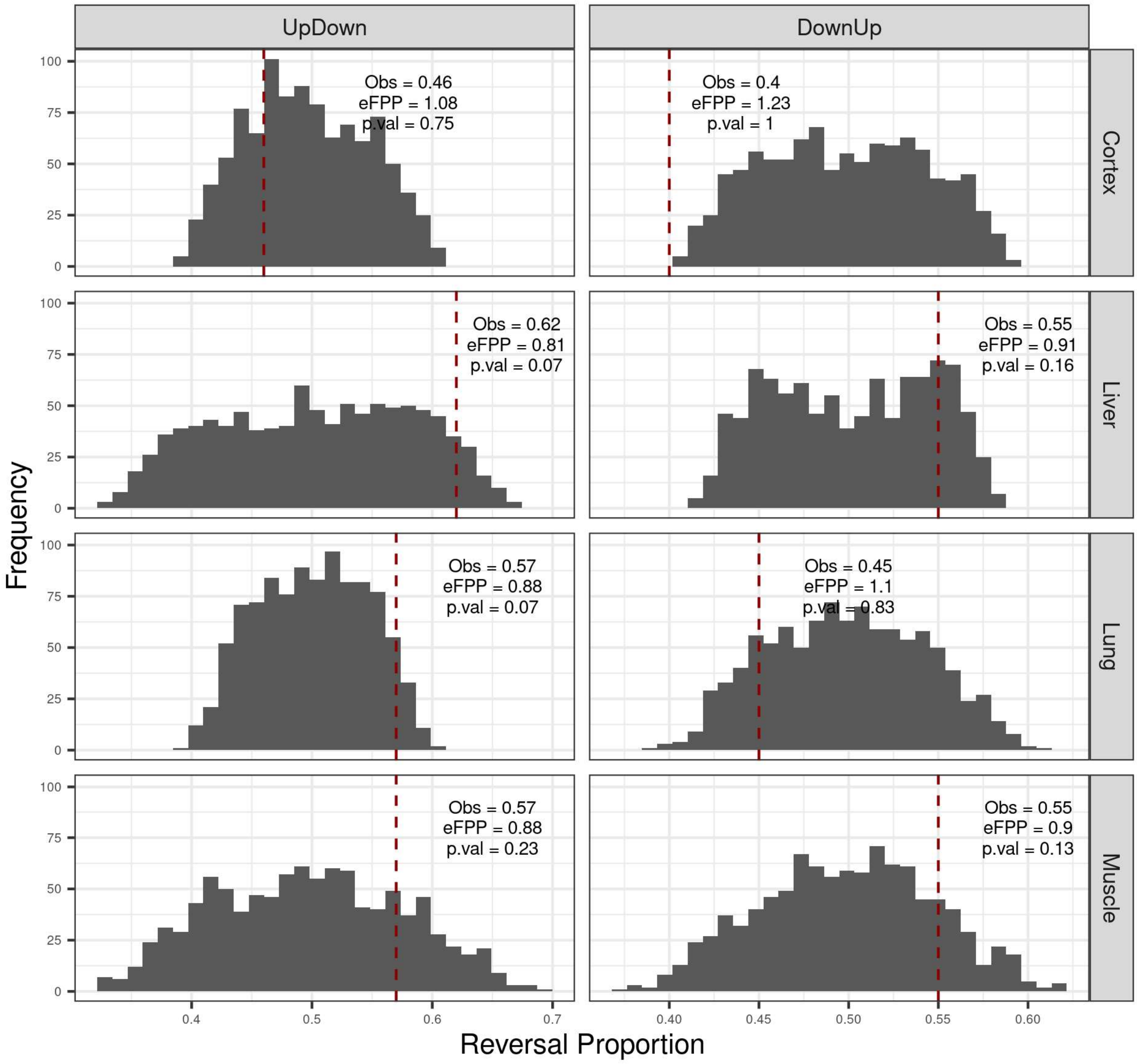


*Permutation test result for up-down and down-up reversal genes in each tissue. Developmental up- (or down-) genes, i.e. genes with expression-age $\rho$>0 (or $\rho$<0), were kept constant and the age labels of the individuals in the ageing period were permuted (Methods). No significance cutoff was used in choosing genes. The dashed red line shows the observed ("Obs") up-down (or down-up) proportions in tissues and eFPP was calculated as the median expected value of the permutations divided by the observed value. P-values were calculated as the proportion of permutations that are higher than or equal to the observed value. Left panel: up-down reversal proportions were calculated as UD/(UD + UU). Right panel: down-up reversal proportions were calculated as DU/(DU+DD).*

**Figure 1-figure supplement 9. Permutation test results for shared reversals among tissues**

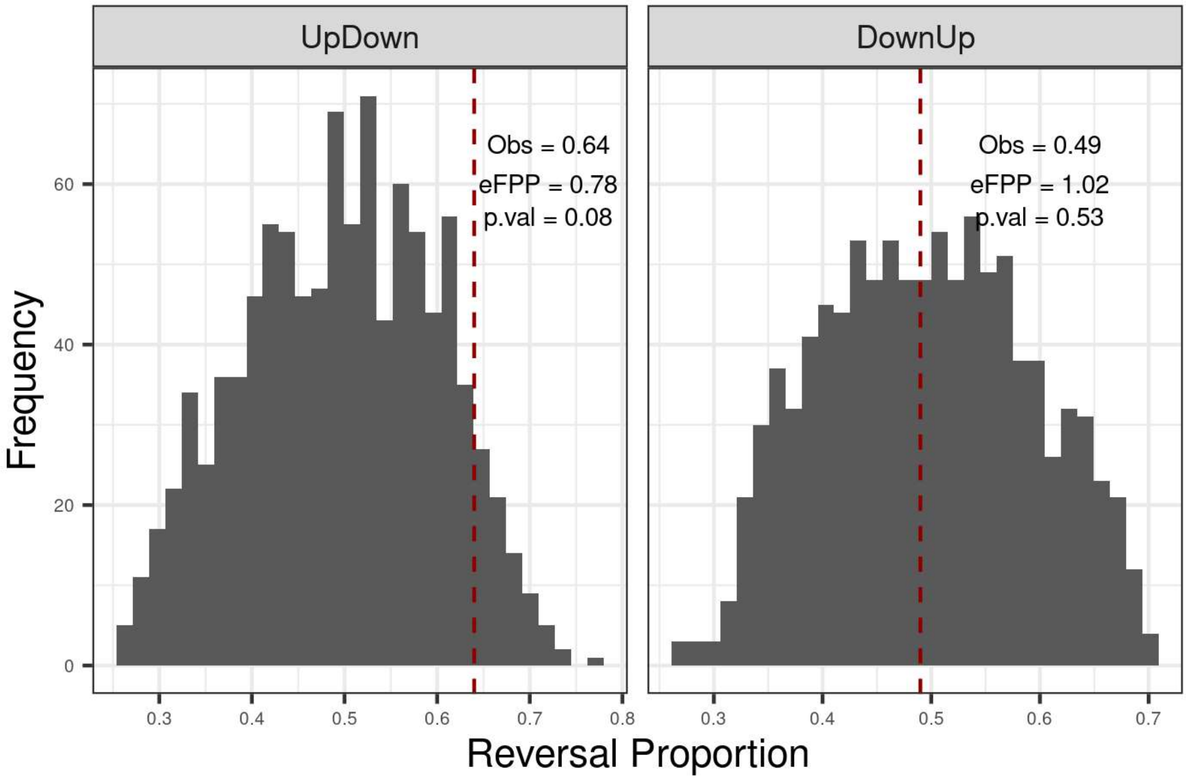


*Permutation test result for shared up-down (or down-up) reversal genes across tissues. Developmental up- (or down-) genes were kept constant (among 2255 shared up-genes and 2209 shared down-genes in development), and the age labels of the individuals in the ageing period were permuted (Methods). The dashed red line shows the observed ("Obs") up-down (or down-up) proportions shared among tissues and eFPP was calculated as the median of the permutations divided by the observed value. The p-values were calculated as the proportion of permutations that are higher than or equal to the observed value. Left panel: up-down reversal proportions were calculated as UD/(UD + UU). Right panel: down-up reversal proportions were calculated as DU/(DU+DD).*

**Figure 1-figure supplement 10. Replication of Figure 1 results using VST normalisation**

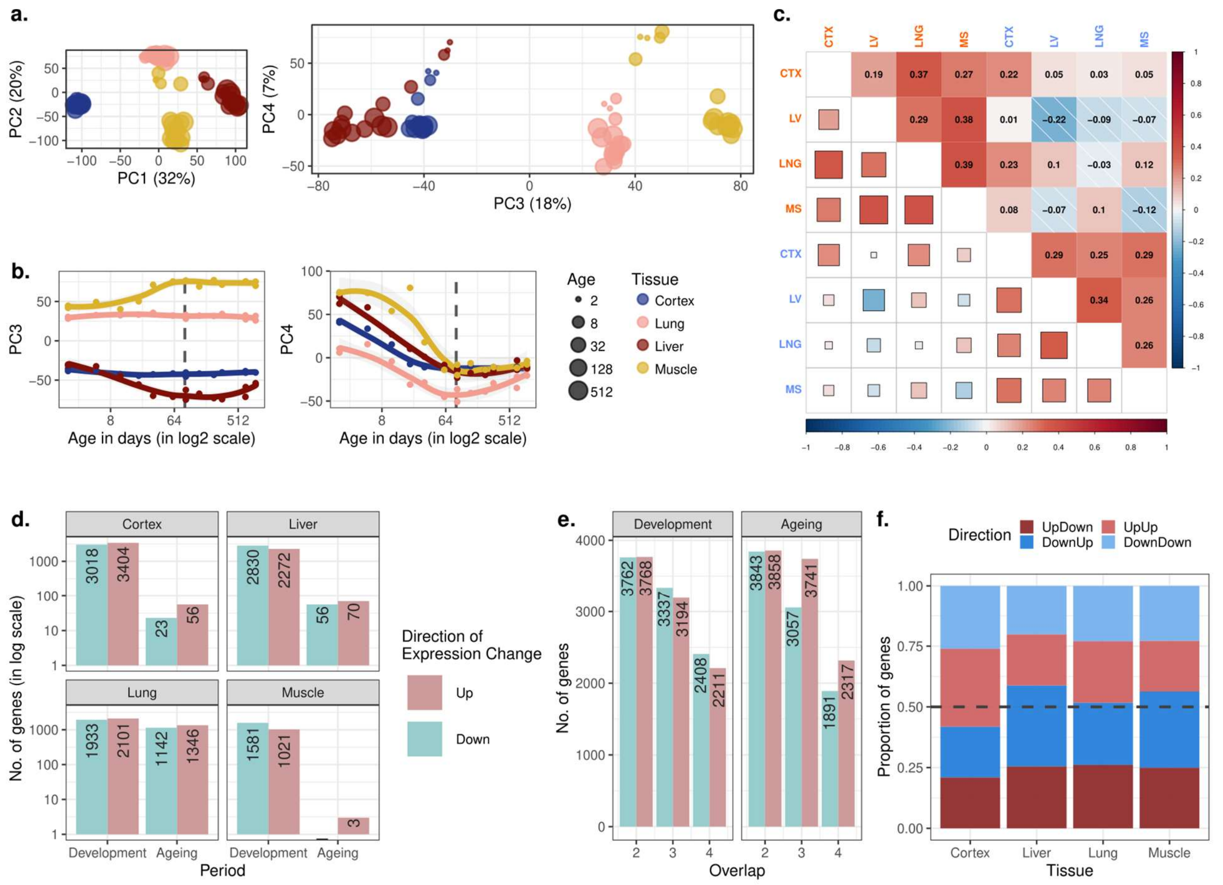


*To confirm the robustness of the results to the choice of normalisation method, the analysis was repeated using an alternative normalisation approach, VST, implemented in the DESeq2 package (see Methods).* ***a)*** *Principal components analysis (PCA) of expression levels of 14,973 protein-coding genes across four tissues of 16 mice. Values in parentheses show the variation explained by each component.* ***b)*** *Age trajectories of PC3 (left) and PC4 (right). Spearman's correlation coefficients between PC4 and age in each tissue in development range between 0.58 and 0.99 (See* ***Figure 1-source data*** *for all tests). The dashed vertical line indicates 90 days of age, separating development and ageing periods.* ***c)*** *Similarity between the age-related gene expression changes (Spearman's correlation coefficient between expression and age without a significance cutoff) across tissues in development and ageing. Similarities were calculated using Spearman's correlation coefficient between expression-age correlations across tissues. CTX: cortex, LV: liver, LNG: lung, MS: muscle.* ***d)*** *The number of significant age-related genes in each tissue (FDR corrected p-value<0.1).* ***e)*** *Shared age-related genes among tissues identified without using a significance cutoff. The x-axis shows the number of tissues among which age-related genes are shared.* ***f)*** *The proportion of age-related expression change trends in each tissue across the lifetime. No significance cutoff was used. UpDown: up-regulation in development and*

*down-regulation in the ageing; DownUp: down-regulation in development and up-regulation in the ageing; UpUp: up-regulation in development and up-regulation in the ageing; DownDown: down-regulation in development and down-regulation in ageing.*

**Figure 1-figure supplement 11. Correlation between QN and VST normalisation methods using age-related expression changes**

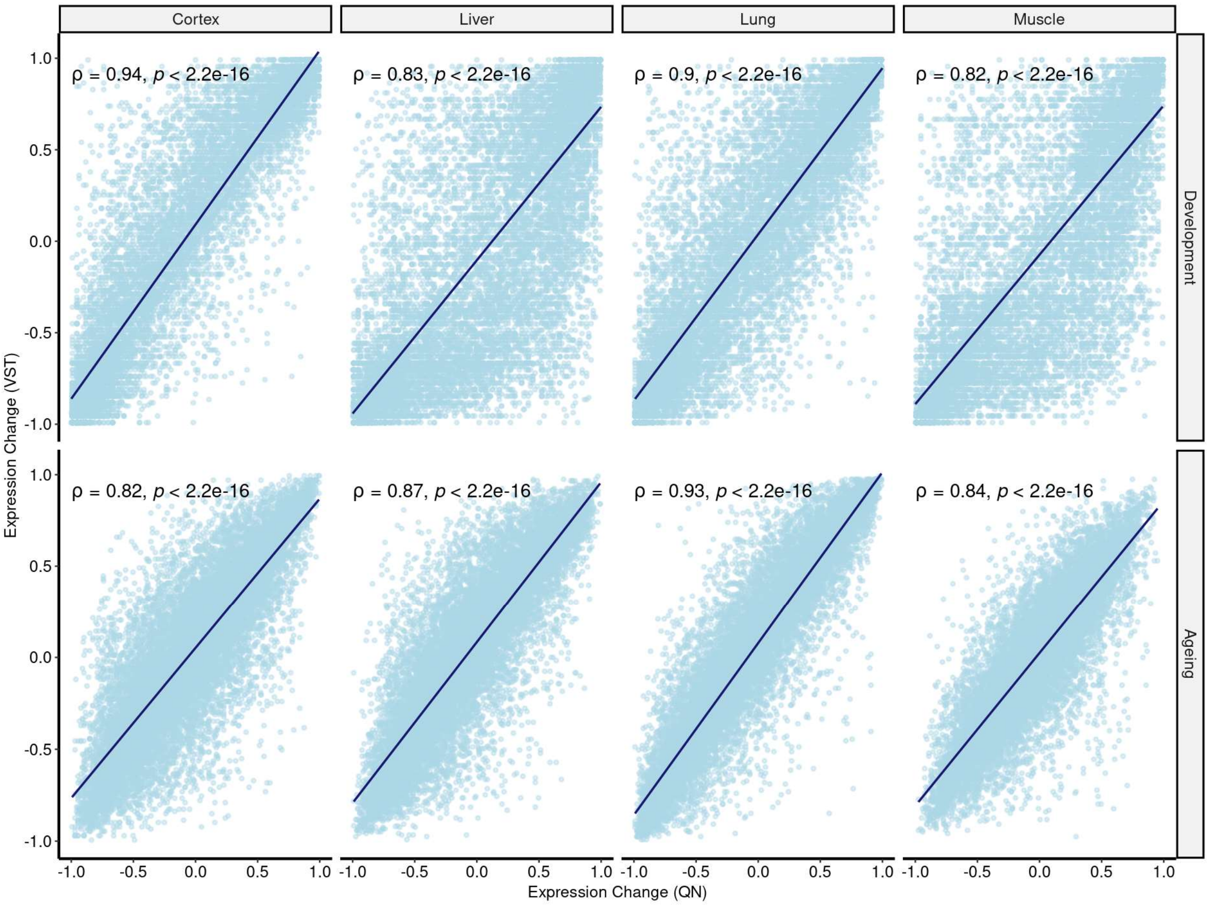


*Spearman's correlation coefficient between expression trajectories of QN (quantile normalised, x-axis) and VST (variance stabilising transformation method from DESeq2 package, y-axis) normalised data. Expression trajectories were calculated using Spearman's correlation coefficient between age and expression level for each gene in both periods ($n_{dev}$ = [14705, 14710], $n_{ageing}$ = [14689, 14710]). Blue lines represent the regression lines.*

**Figure 1-figure supplement 12. Clustering of genes by expression levels in cortex tissue**

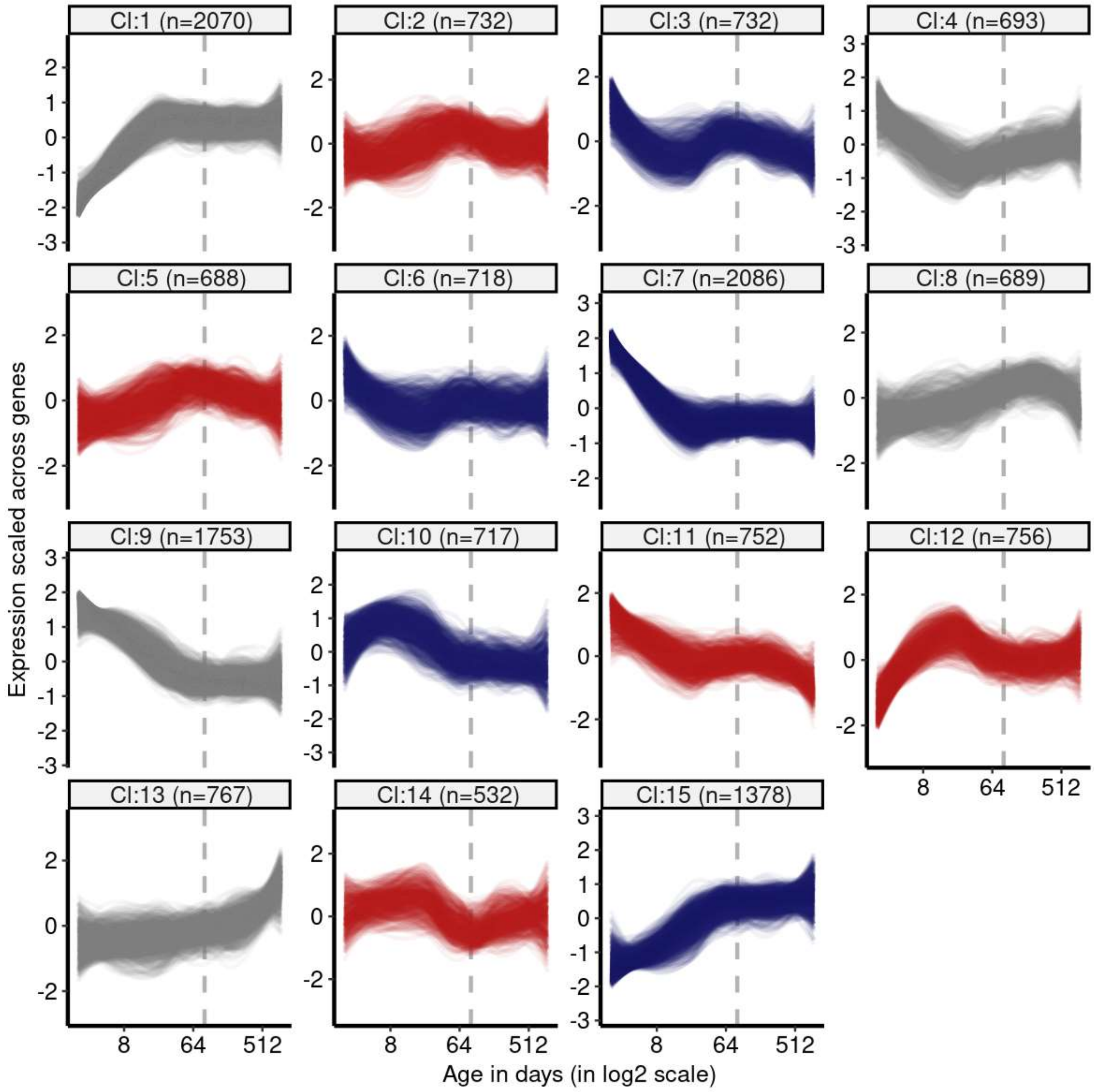


*K-means clustering (k=15) of genes (15,063) using expression levels in cortex tissue. Numbers in the parentheses show the number of genes in each cluster. Expression levels of genes were scaled across samples (mean=1, sd=0) before clustering. The optimal number of clusters was determined with gap statistics (see Methods). Clusters enriched among DiCo genes compared to all other clusters were indicated with red colour and the ones depleted among DiCo genes were indicated with blue colour. The list of genes belonging to each cluster and their enrichment among DiCo genes are given in **Figure 1-source data**.*

**Figure 1-figure supplement 13. Clustering of genes by expression levels in lung tissue**

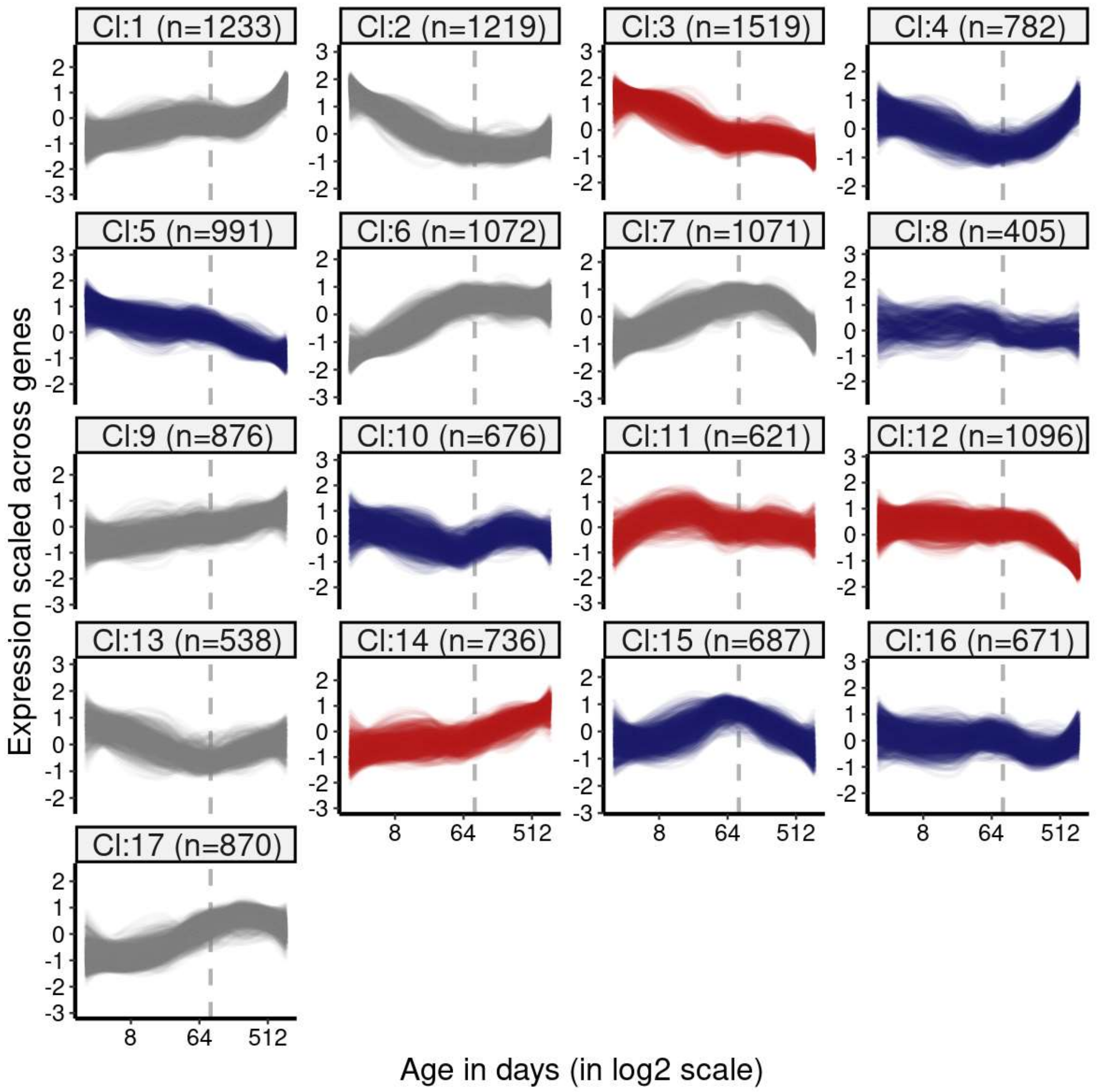


*K-means clustering (k=17) of genes (15,063) using expression levels in lung tissue. Numbers in the parentheses show the number of genes in each cluster. Expression levels of genes were scaled across samples (mean=1, sd=0) before clustering. The optimal number of clusters was determined with gap statistics (See Methods). Clusters enriched among DiCo genes were indicated with red colour and the ones depleted among DiCo genes were indicated with blue colour. The list of genes belonging to each cluster and their enrichment among DiCo genes are given in* ***Figure 1-source data****.*

**Figure 1-figure supplement 14. Clustering of genes by expression levels in liver tissue**

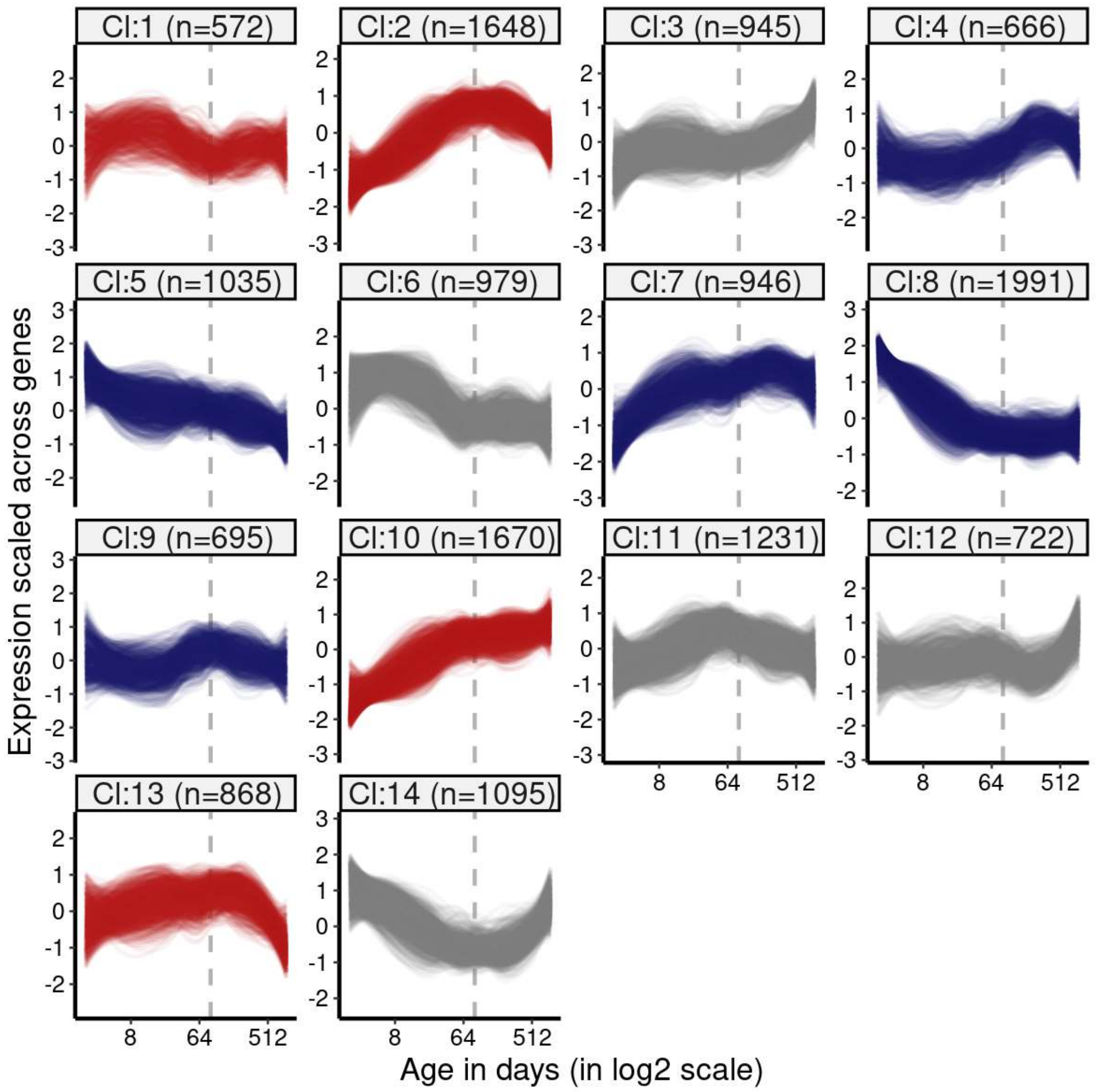


*K-means clustering (k=14) of genes (15,063) using expression levels in liver tissue. Numbers in the parentheses show the number of genes in each cluster. Expression levels of genes were scaled across samples (mean=1, sd=0) before clustering. The optimal number of clusters was determined with gap statistics (See Methods). Clusters enriched among DiCo genes were indicated with red colour and the ones depleted among DiCo genes were indicated with blue colour. The list of genes belonging to each cluster and their enrichment among DiCo genes are given in* ***Figure 1-source data****.*

**Figure 1-figure supplement 15. Clustering of genes by expression levels in muscle tissue**

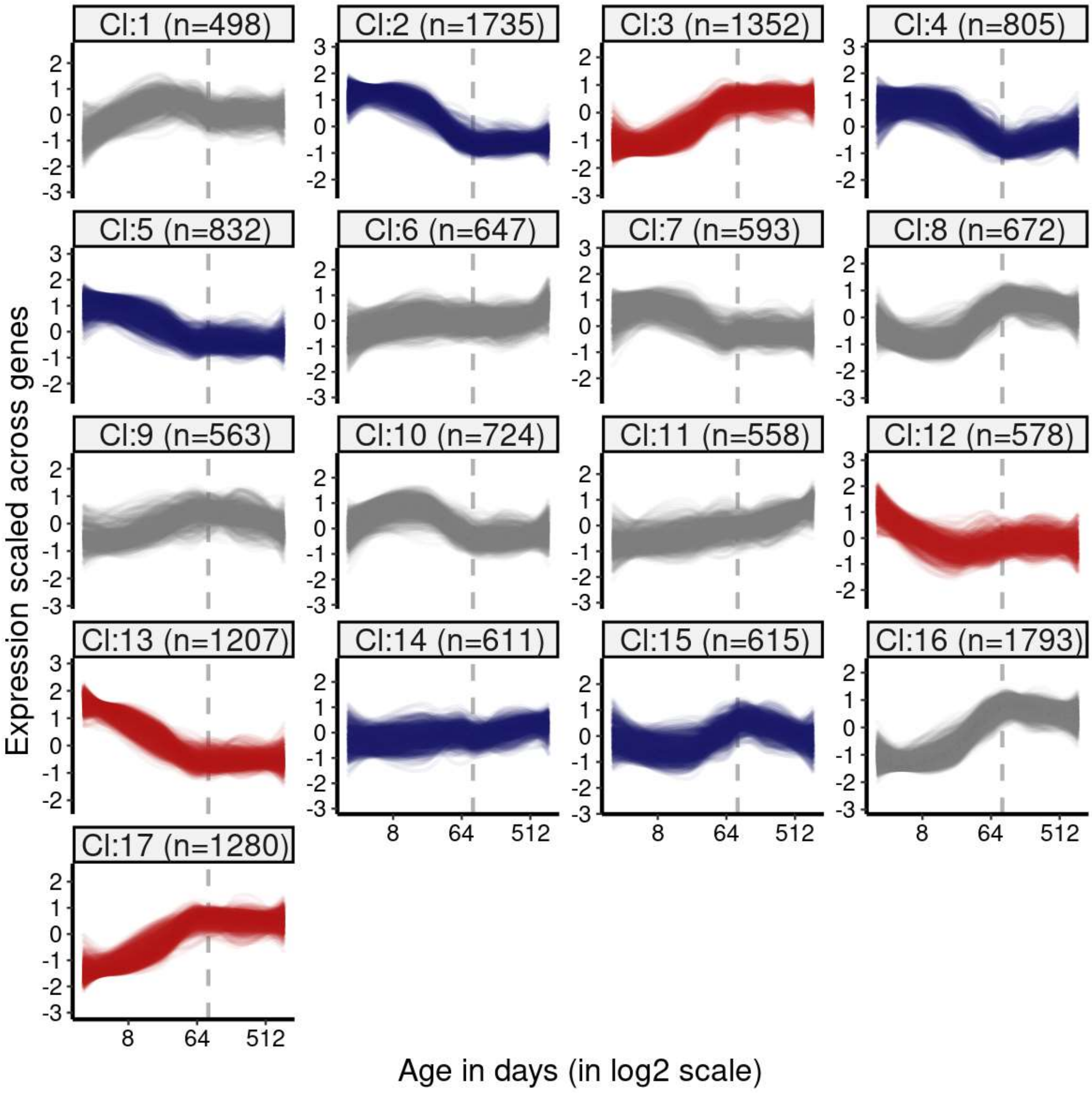


*K-means clustering (k=17) of genes (15,063) using expression levels in muscle tissue. Numbers in the parentheses show the number of genes in each cluster. Expression levels of genes were scaled across samples (mean=1, sd=0) before clustering. The optimal number of clusters was determined with gap statistics (See Methods). Clusters enriched among DiCo genes were indicated with red colour and the ones depleted among DiCo genes were indicated with blue colour. The list of genes belonging to each cluster and their enrichment among DiCo genes are given in* ***Figure 1-source data****.*

**Figure 2-figure supplement 1. Age-related change in CoV summarised across genes using median CoV values**

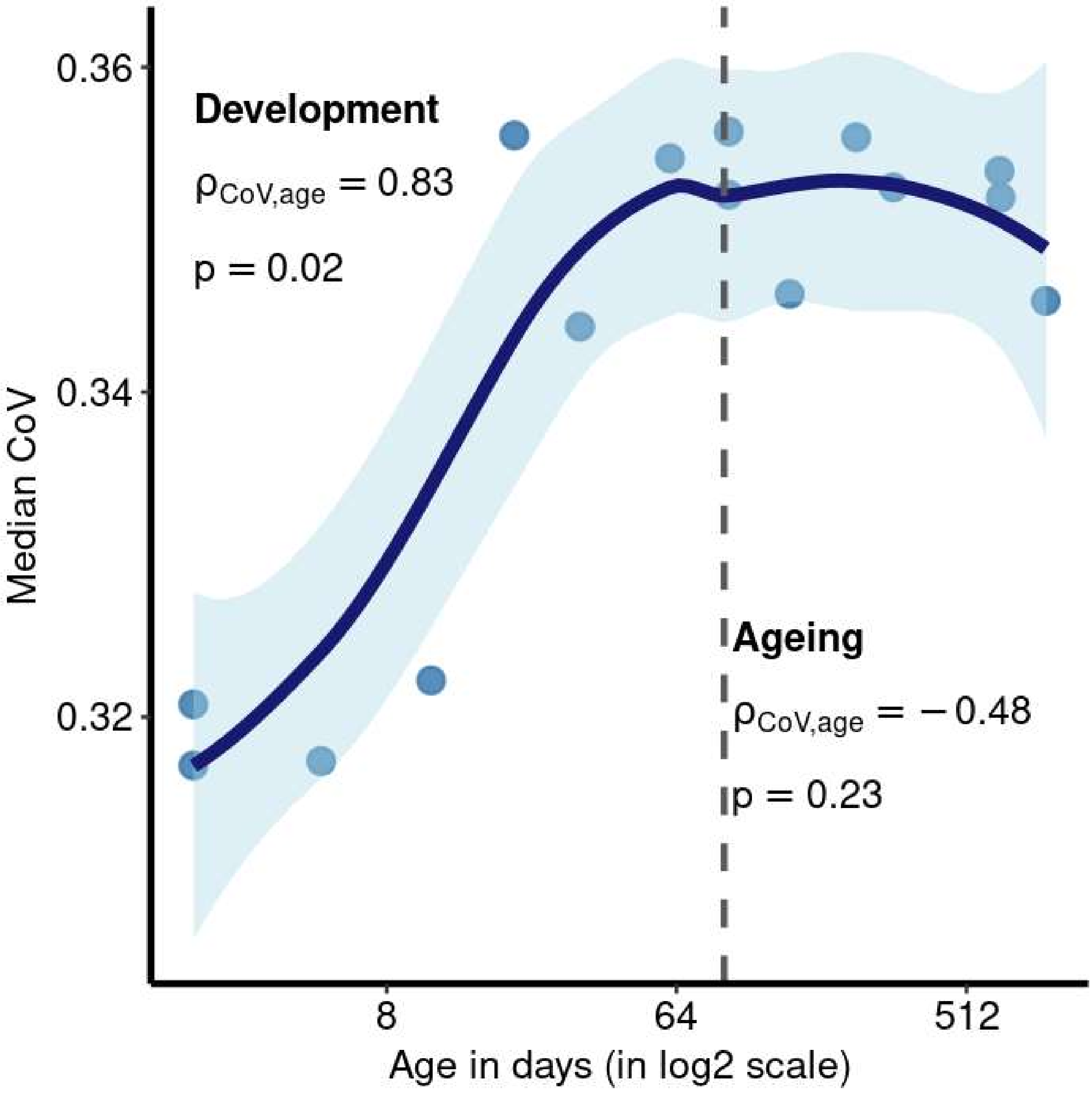


*Each point represents the median CoV value (instead of the mean given in Figure 2a) of all protein-coding genes (15,063) for each mouse except the one that lacks expression data in the cortex (n=15). x-axis is in log2 scale. The dashed grey line shows the start of the ageing period. The Spearman's correlation coefficient and p-value for each period are indicated separately on the plot.*

**Figure 2-figure supplement 2. Clustering of DiCo genes by expression variations (CoV) among tissues**

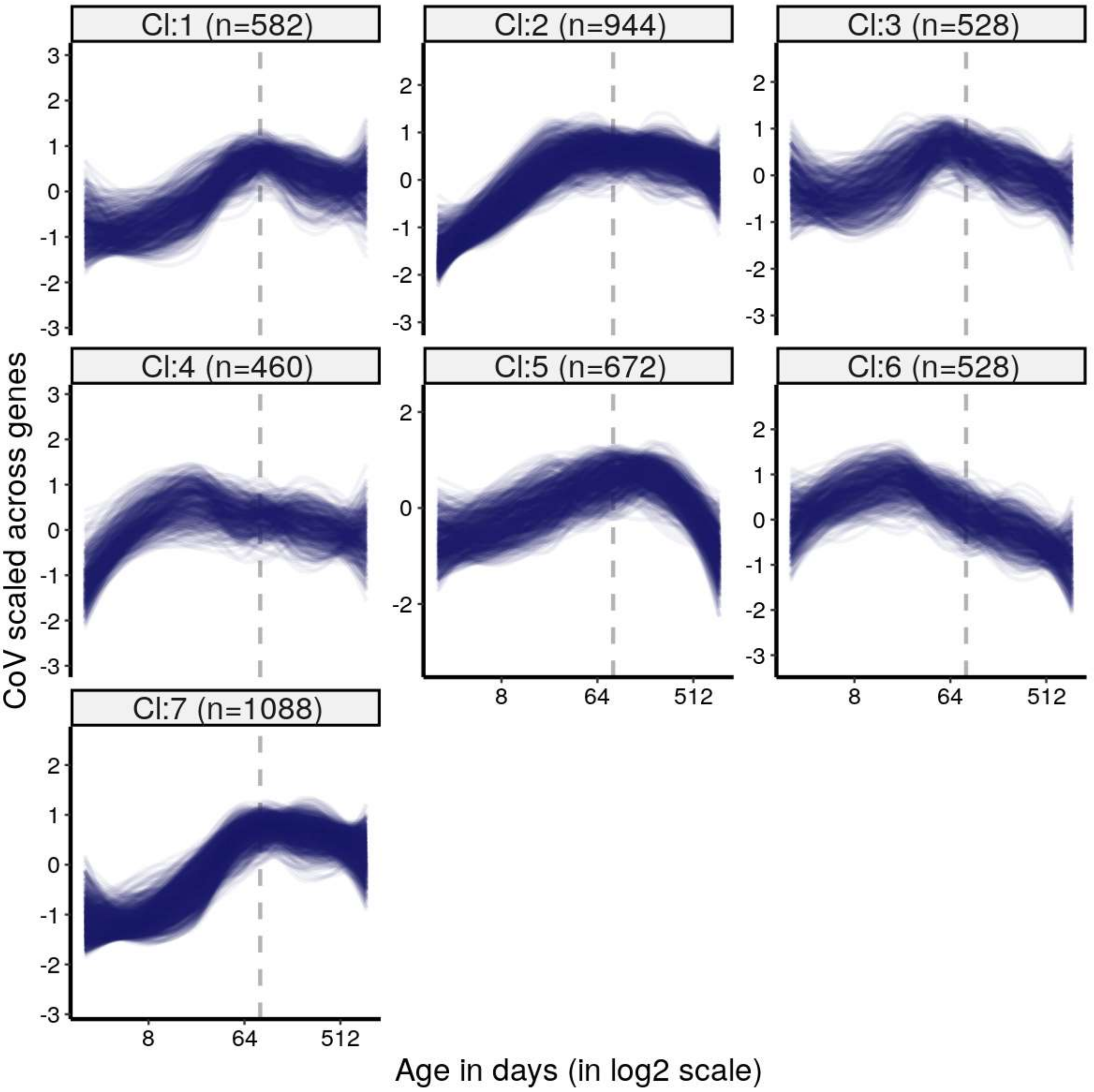


*Kmeans clustering (k=7) of DiCo genes (4,802) using CoV values. Numbers in the parentheses show the number of genes in each cluster. CoV values were scaled across genes (mean=1, sd=0) before clustering. The optimal number of clusters was determined with gap statistics (Methods). The list of genes belonging to each cluster and their age-related CoV change correlations are given in* ***Figure 2-source data****.*

**Figure 2-figure supplement 3. Clustering of DiCo genes by expression levels in tissues**

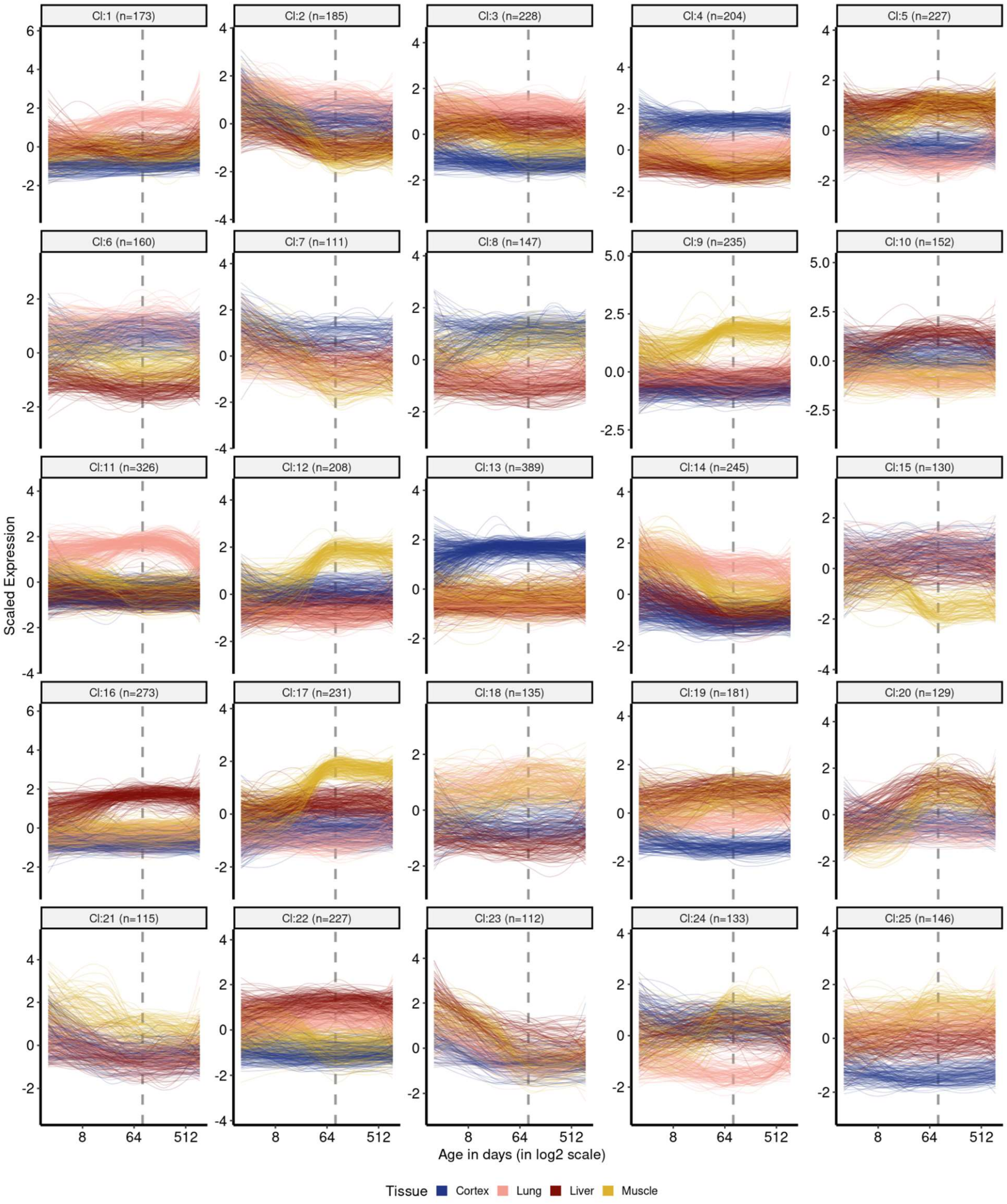


*Kmeans clustering (k=25) of DiCo genes (n=4,802) using gene expression levels. Numbers in the parentheses show the number of genes in each cluster. Expression levels of genes were scaled across tissues ((mean=1, sd=0)) before clustering. The optimal number of clusters was determined with gap statistics (Methods). The list of genes belonging*

*to each cluster and their age-related CoV change correlations are given in **Figure 2-source data**.*

**Figure 2-figure supplement 4. Number of genes with inter-tissue divergence and convergence tendencies in development and ageing**

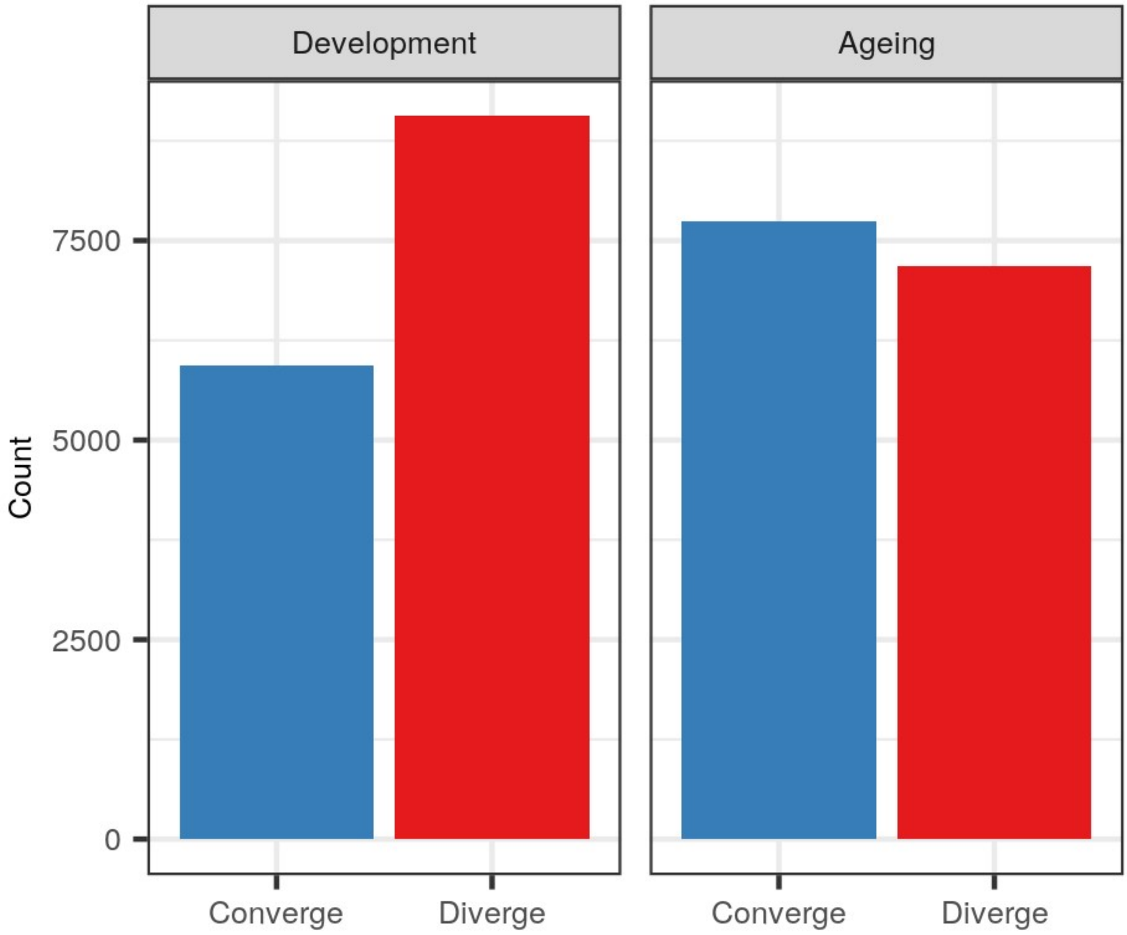


*The number of CoV changes with age (without a significance cutoff) during development and ageing. Converge: genes showing negative correlation ($\rho$<0) between CoV and age; Diverge: genes showing positive correlation ($\rho$>0) between CoV and age (Development: $n_{converge}$=5,939, $n_{diverge}$=9,058; Ageing: $n_{converge}$=7,748, $n_{diverge}$=7,187).*

**Figure 2-figure supplement 5. Pairwise tissue expression correlations**

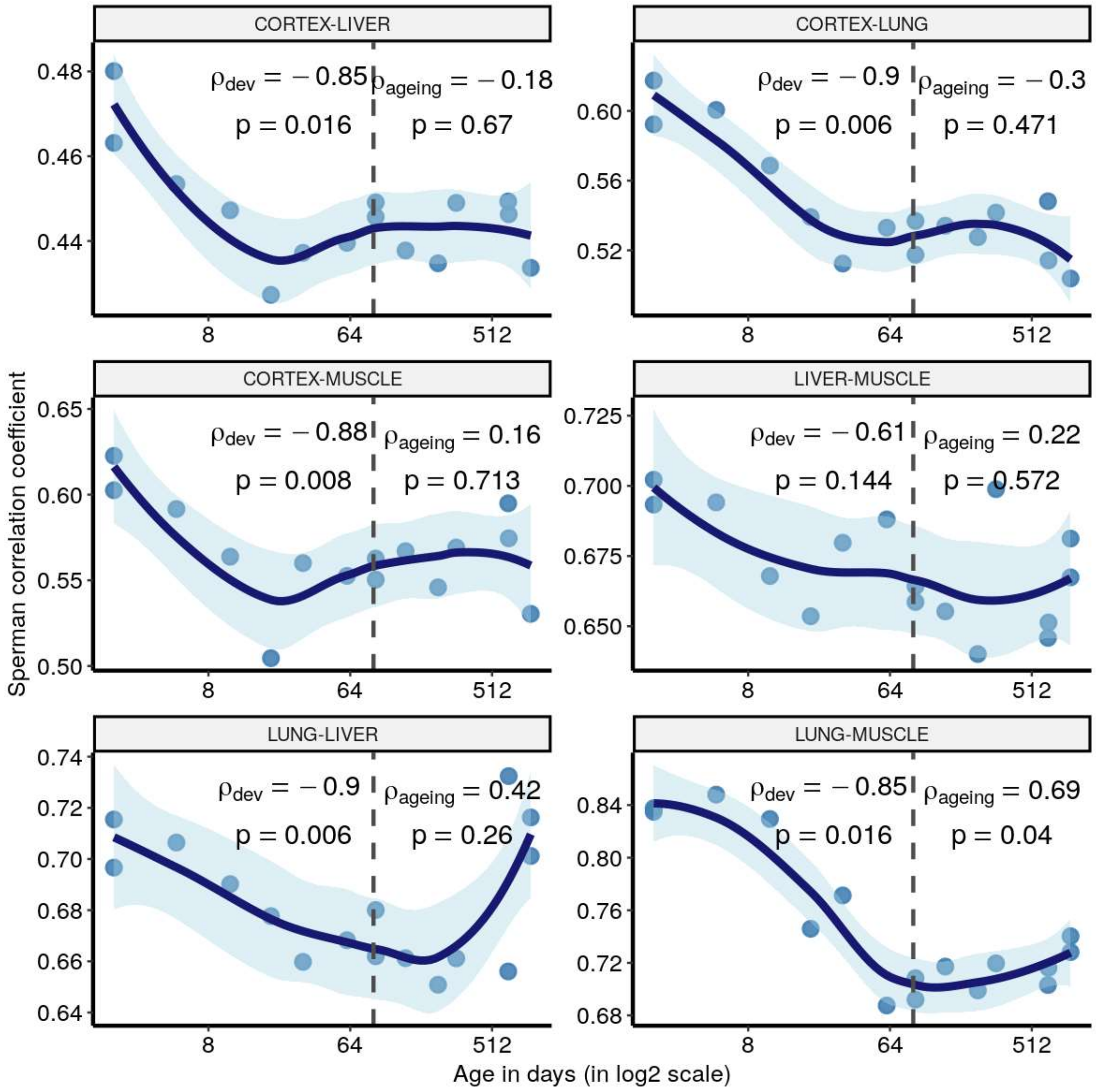


*Age-related changes in pairwise Spearman's correlation coefficients for the expression levels (y-axis) between tissues of the same individual mouse in our dataset. The dashed grey line indicates the start of the ageing period. The Spearman's correlation coefficients and p values for each period are indicated separately on the plot.*

**Figure 2-figure supplement 6. Summary of pairwise expression correlations among tissues**

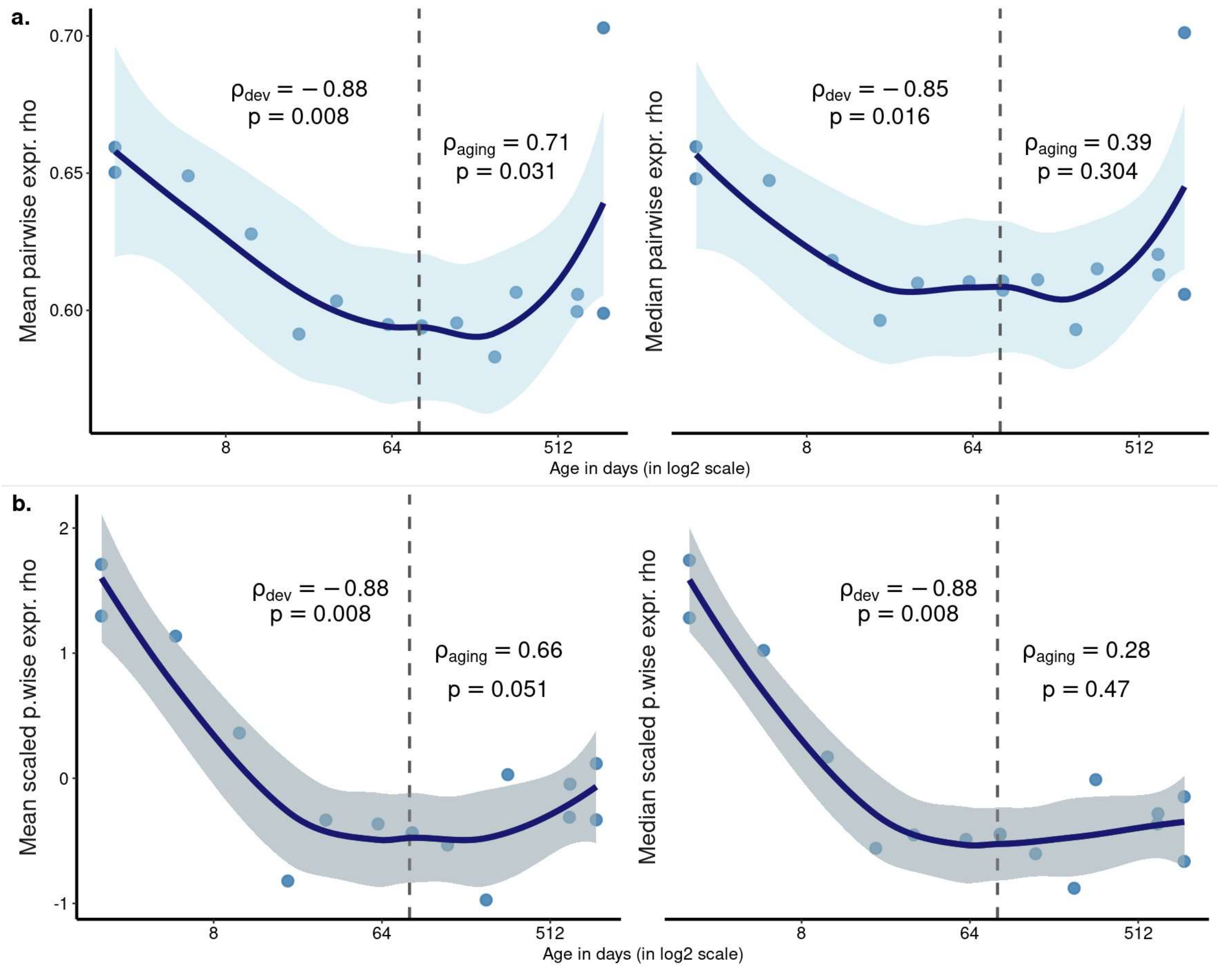


*Age-related change in the mean (left) or the median (right) pairwise expression correlations among tissues. Each point represents the mean (left) or the median (right) of pairwise expression correlations among tissues of the same mouse (mean/median values are calculated from **Figure 2–figure supplement 5**). **a)** Absolute expression correlations were used to calculate the mean or the median. **b)** Expression correlations were scaled within each tissue pair (mean=1, sd=0) before calculating the mean and median. The Spearman's correlation coefficients and p values for each period are indicated separately on the plot.*

**Figure 2-figure supplement 7. CoV and pairwise correlation analysis of Jonker dataset**

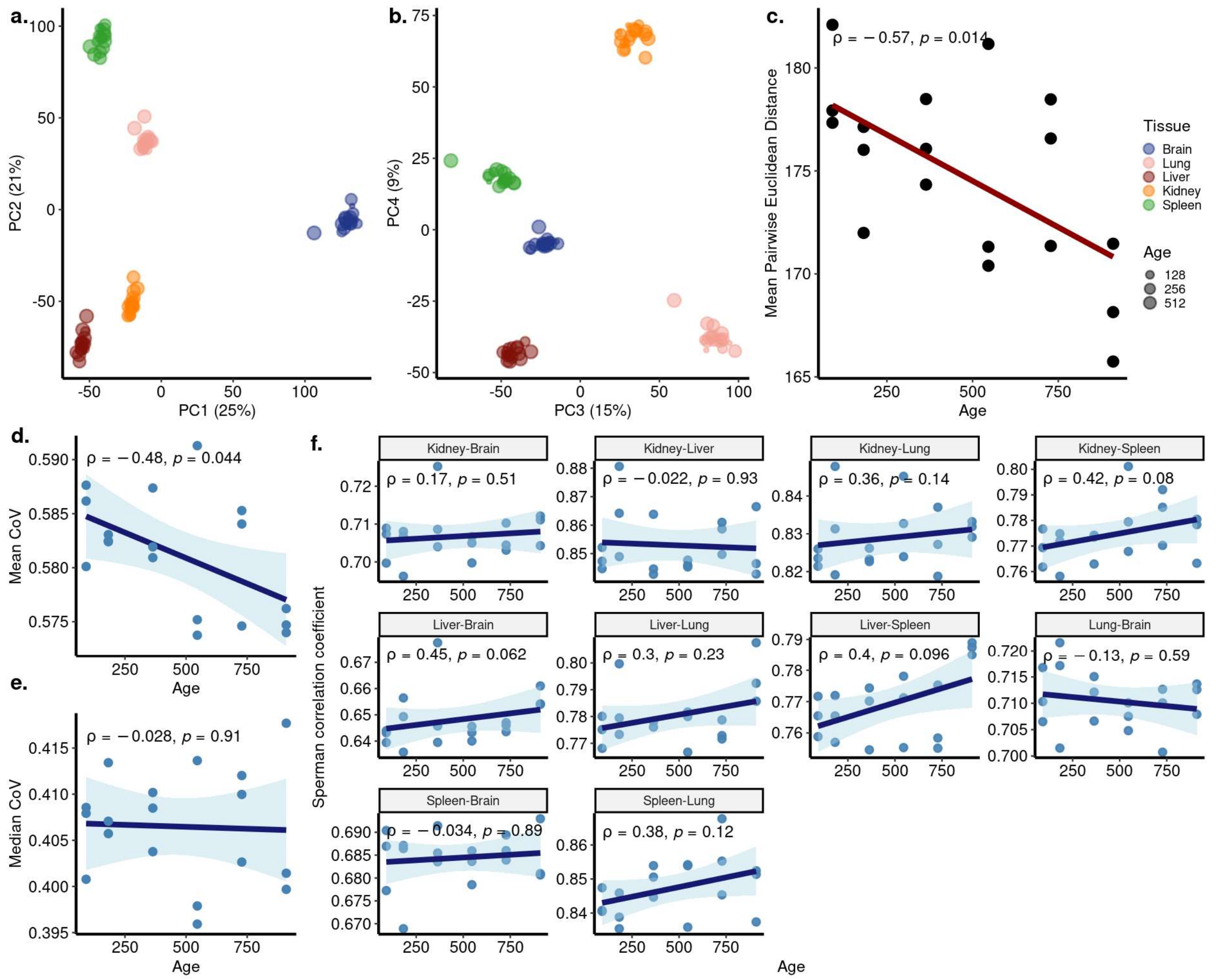


***a-b)** Principal components analysis (PCA) of expression values of 17,661 protein-coding genes across five tissues (Brain (Cortex), Liver, Lung, Kidney, Spleen) of 18 individuals in the Jonker dataset (contains samples only from the ageing period). Values in parentheses show the variance explained by each PC. **c)** The change in mean pairwise Euclidean distance between the PC values for the tissues of the same individuals (y-axis) with age (x-axis). Transcriptome-wide **d)** mean and **e)** median CoV changes with age across 5 tissues. The x-axis shows age in days. Each point represents the mean or median CoV value of all protein-coding genes for each individual. **f)** Spearman's correlation coefficient between age (x-axis) and gene expression correlations of each individual in pairwise tissues (y-axis). Spearman's correlation coefficient and p-values are indicated in each plot.*

**Figure 2-figure supplement 8. PCA of GTEx dataset covering cortex, liver, lung, and muscle tissues**

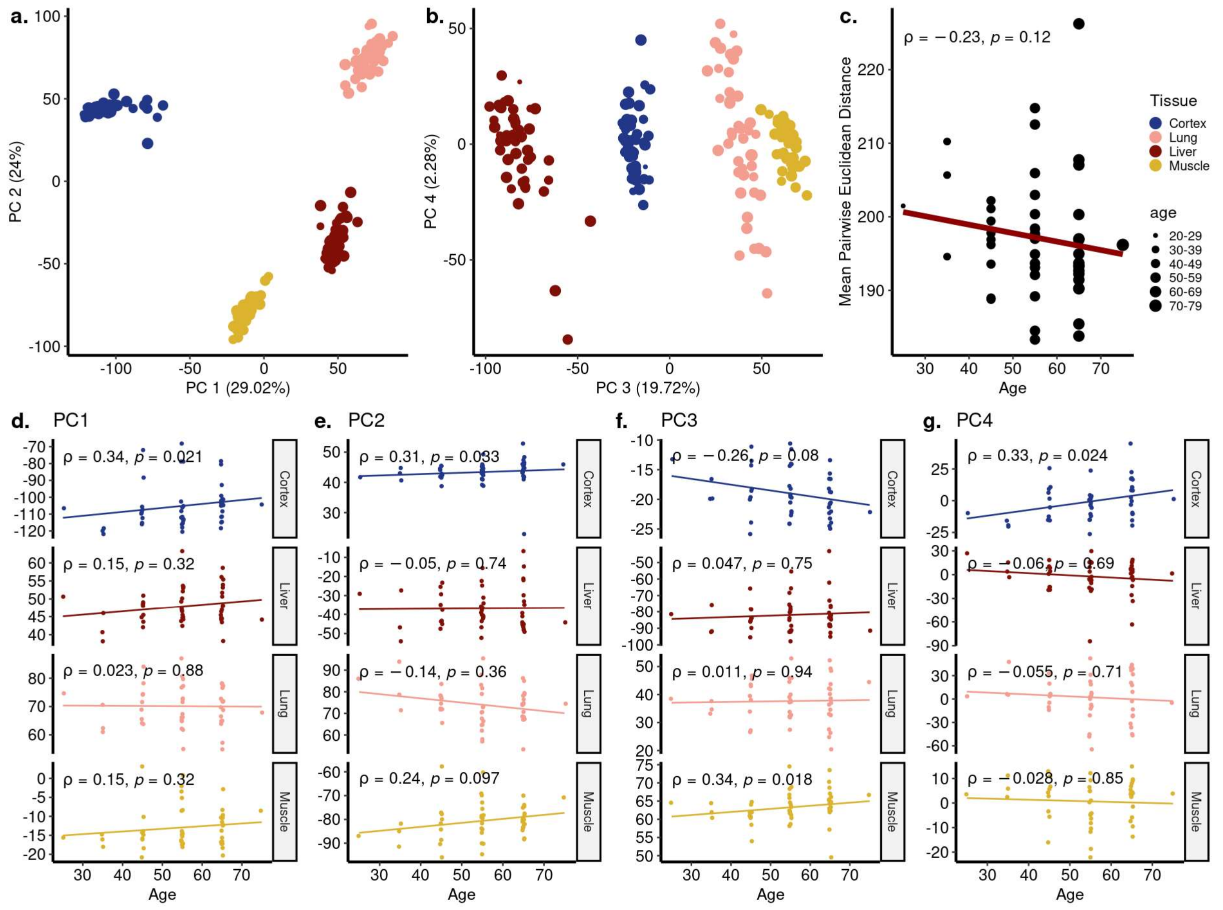


*a-b) Principal components analysis (PCA) of expression values of 16,197 genes across four tissues (Cortex, Liver, Lung, Muscle) of 47 individuals in GTEx. Values in parentheses show the variance explained by each PC. **c)** The change in mean pairwise Euclidean distance between the PC values for the tissues of the same individuals (y-axis) with age (x-axis). **d-g)** Association between the first four PCs (y-axis) and age (x-axis). The tissue and age of the samples are indicated by the colour and size of the points, respectively. Spearman's correlation test results are indicated in each plot.*

**Figure 2-figure supplement 9. CoV and pairwise correlation analysis of GTEx dataset covering cortex, liver, lung, and muscle tissues**

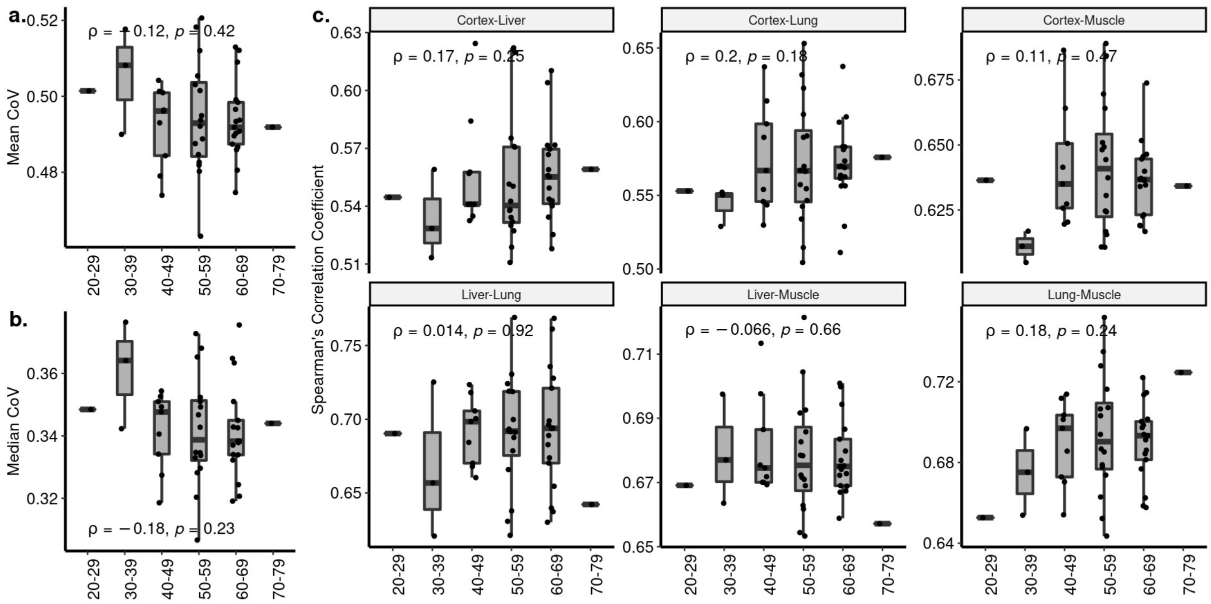


***a-b)** Transcriptome-wide mean **(a)** and median **(b)** CoV change with age across four tissues (Cortex, Liver, Lung, Muscle) in GTEx. Each point represents the mean or median CoV value of all protein-coding genes (16,197) for each individual (n=47) in GTEx. Spearman's correlation coefficients and p-values are also presented in the plot. **c)** The change in pairwise Spearman's correlation coefficient between gene expression values of the same individual across ages (y-axis) with age (x-axis). Spearman's correlation coefficient and p-values between the pairwise tissue correlations and age are also presented in each plot.*

**Figure 2–figure supplement 10. PCA of GTEx dataset with ten tissues**

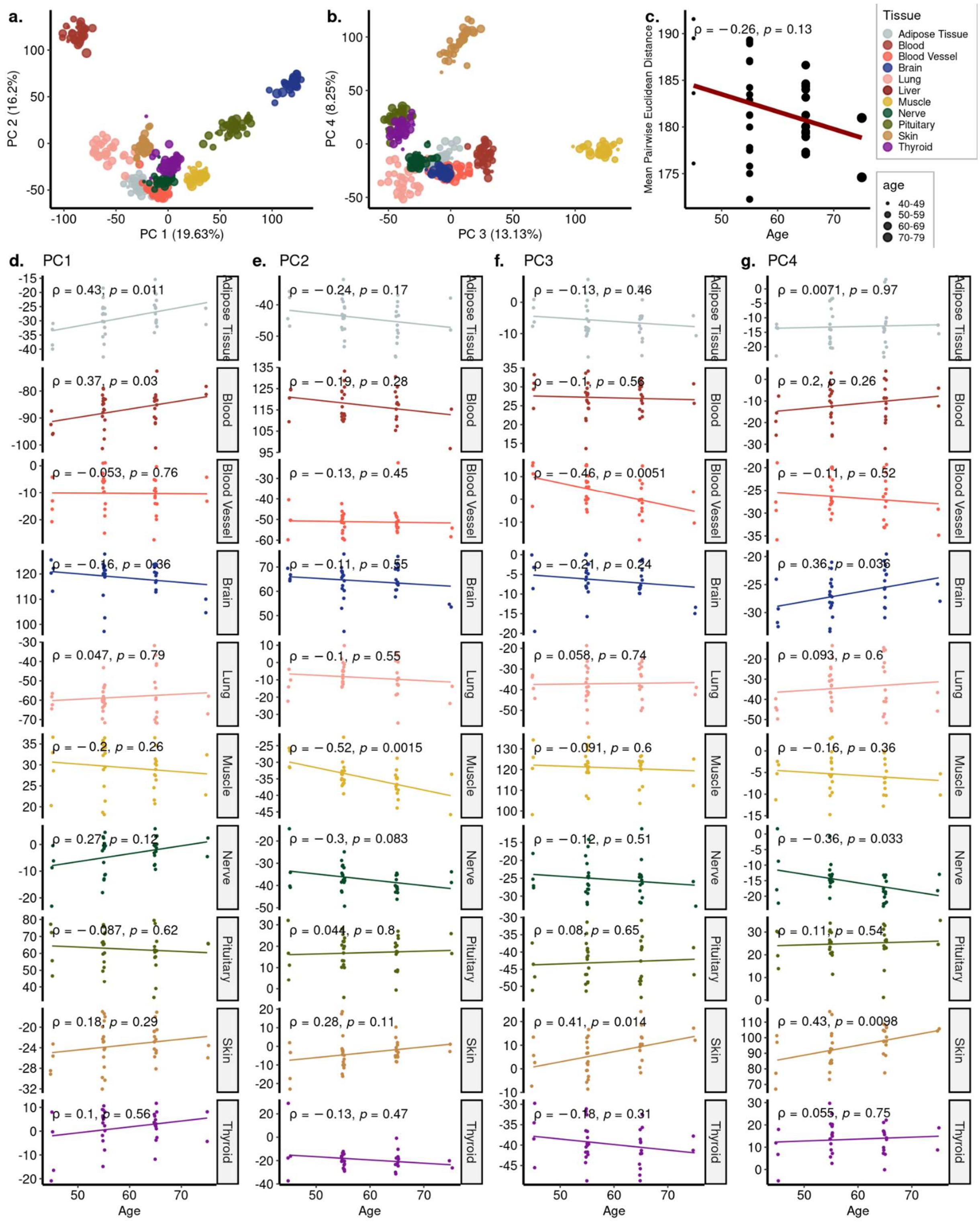

***a-b)*** *Principal components analysis (PCA) of expression values of 16,290 genes across ten tissues of 35 individuals in GTEx. Values in parentheses show the variance explained by each PC.* ***c)*** *The change in mean pairwise Euclidean distance between the PC values for the tissues of the same individuals (y-axis) with age (x-axis).* ***d-g)*** *Association between the first four PCs (y-axis) and age (x-axis). The tissue and age of the samples are indicated by the colour and size of the points, respectively.*

**Figure 2-figure supplement 11. CoV and pairwise correlation analysis of GTEx dataset with ten tissues**

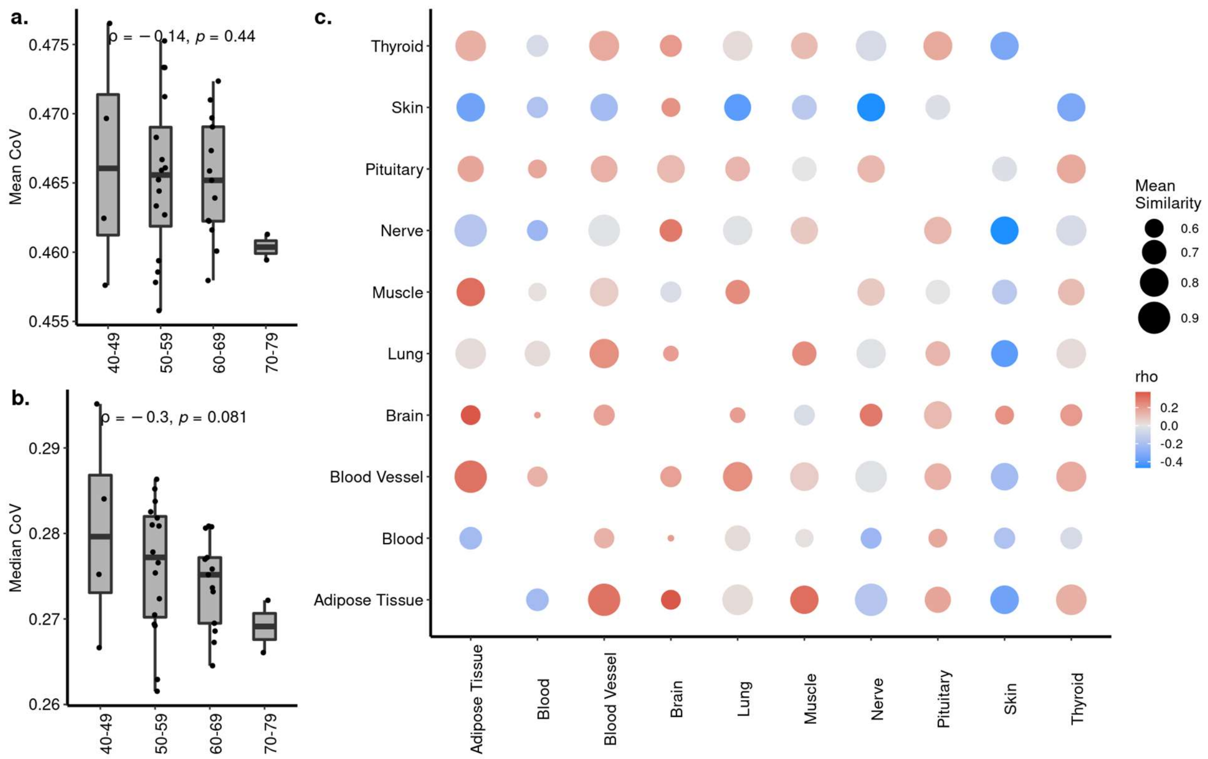


*a-b) Transcriptome-wide mean (a) and median (b) CoV change with age across ten tissues in GTEx. Each point represents the mean or median CoV value of all protein-coding genes (16,290) for each individual (n=35) in GTEx. Spearman's correlation coefficients and p-values are also presented in the plot. c) Age-related changes in pairwise Spearman's correlation coefficient between gene expression values of the same individual. The colour of points shows the correlations between age and pairwise correlations, where darker red colour indicates an increased correlation with age and darker blue indicates a decreased correlation. The size of points shows the mean similarity (correlation) between tissues using all ages. None of the correlations is significant after multiple testing correction (using BH).*

**Figure 2-figure supplement 12. Permutation test result for the proportion of DiCo genes**

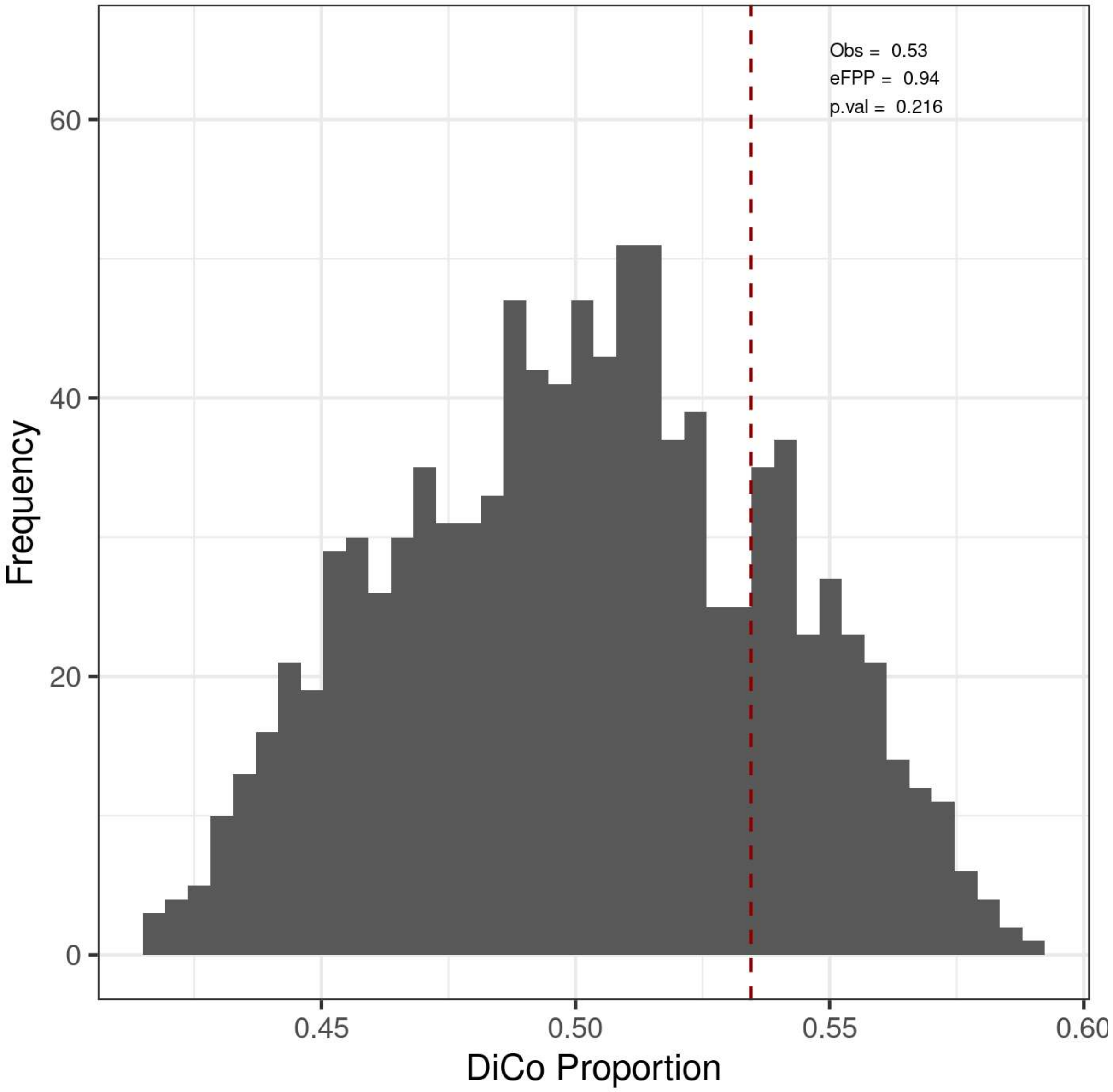


*DiCo genes (n=4,802) were tested with a permutation-based test explained in Methods. We kept the divergent genes (n=9,058) in development constant and permuted age labels of individuals in the ageing period. Then, we calculated the DiCo proportion among those genes in permutations. "Obs:" observed DiCo proportion (Obs = 4,802/9,058, i.e. DiCo/(DiCo + Di~); Di~: divergence across lifetime). eFPP was calculated as the median expected proportion divided by the observed value. P-value was calculated as the proportion of permutations that are higher than or equal to the observed value.*

**Figure 2-figure supplement 13. Clustering of tissues by the presence of samples from the same individuals**

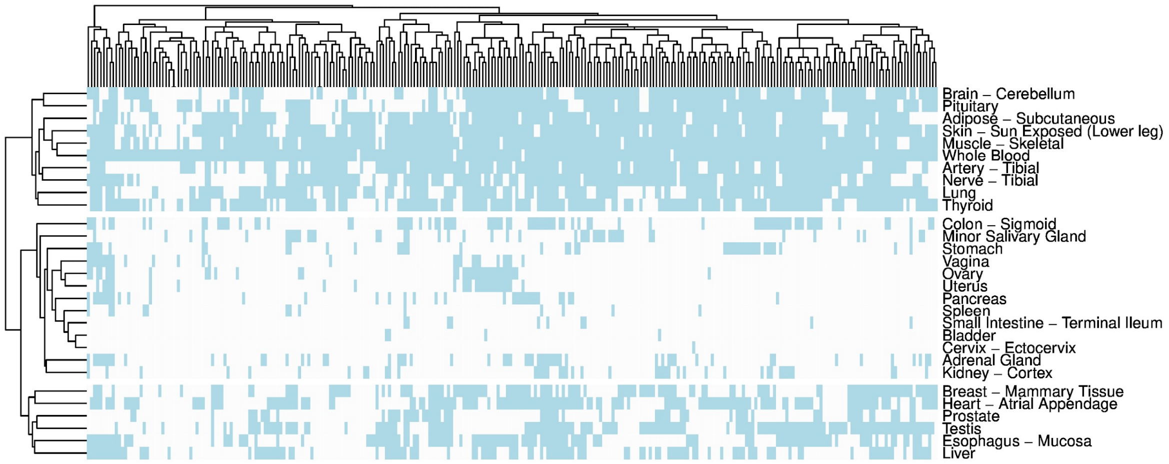


*Heatmap showing whether individuals (columns) have samples (light blue colour) in tissues (y-axis).*

**Figure 2-figure supplement 14. Reproducing Figure 2 results with VST normalisation**

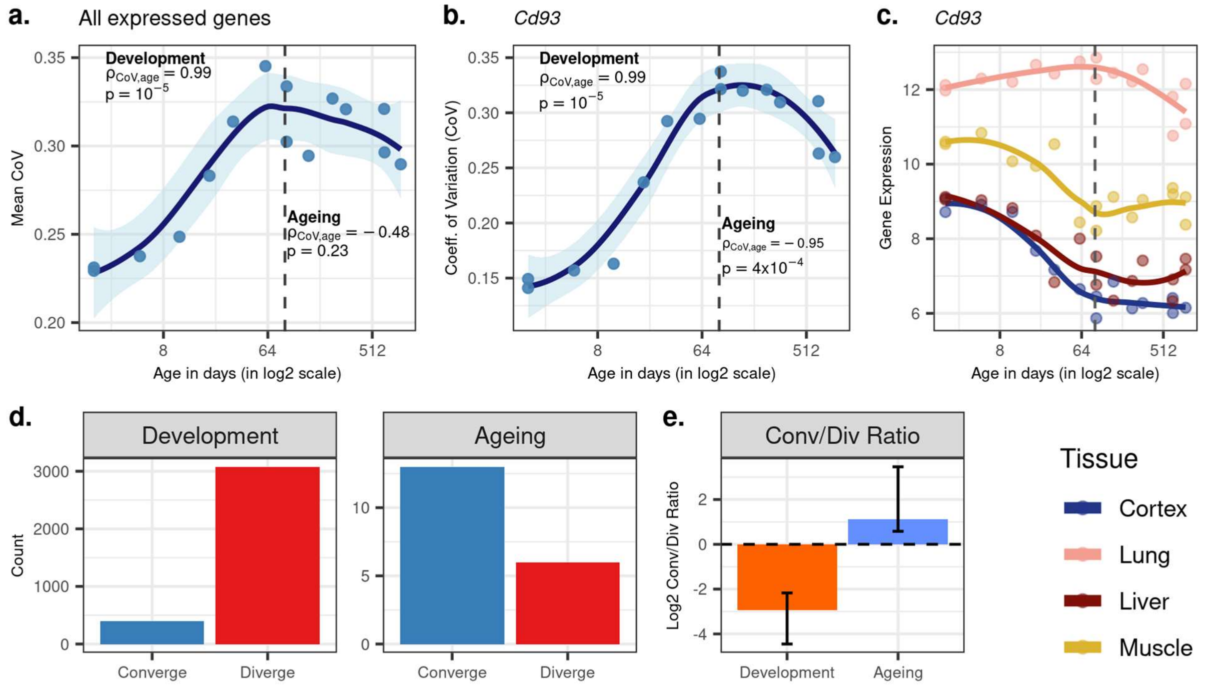


***a)** Transcriptome-wide mean CoV trajectory with age. Each point represents the mean CoV value of all protein-coding genes (14,973) for each mouse (n=15) except the one that lacks expression data in the cortex. **b)** Age effect on CoV value of the Cd93 gene which has the highest rank for the DiCo pattern, in four tissues (Methods). CoV increases during development and decreases during ageing, indicating expression levels show DiCo patterns among tissues. **c)** Expression trajectories of the gene Cd93 in four tissues. **d)** The number of significant CoV changes with age (FDR corrected p-value <0.1) during development (left, $n_{conv.}$=398, $n_{div.}$=3,078) and ageing (right, $n_{conv.}$=13, $n_{div.}$=6). Converge: genes showing a negative correlation (ρ) between CoV and age; Diverge: genes showing a positive correlation between CoV and age. **e)** Log2 ratio of convergent/divergent genes in development and in ageing. The graph represents only genes showing significant CoV changes (at FDR corrected p-value <0.1, given in panel d). Error bars represent the range of log2 ratios calculated from leave-one-out samples in jackknife procedure.*

**Figure 2-figure supplement 15. Effect of heteroscedasticity to DiCo pattern**

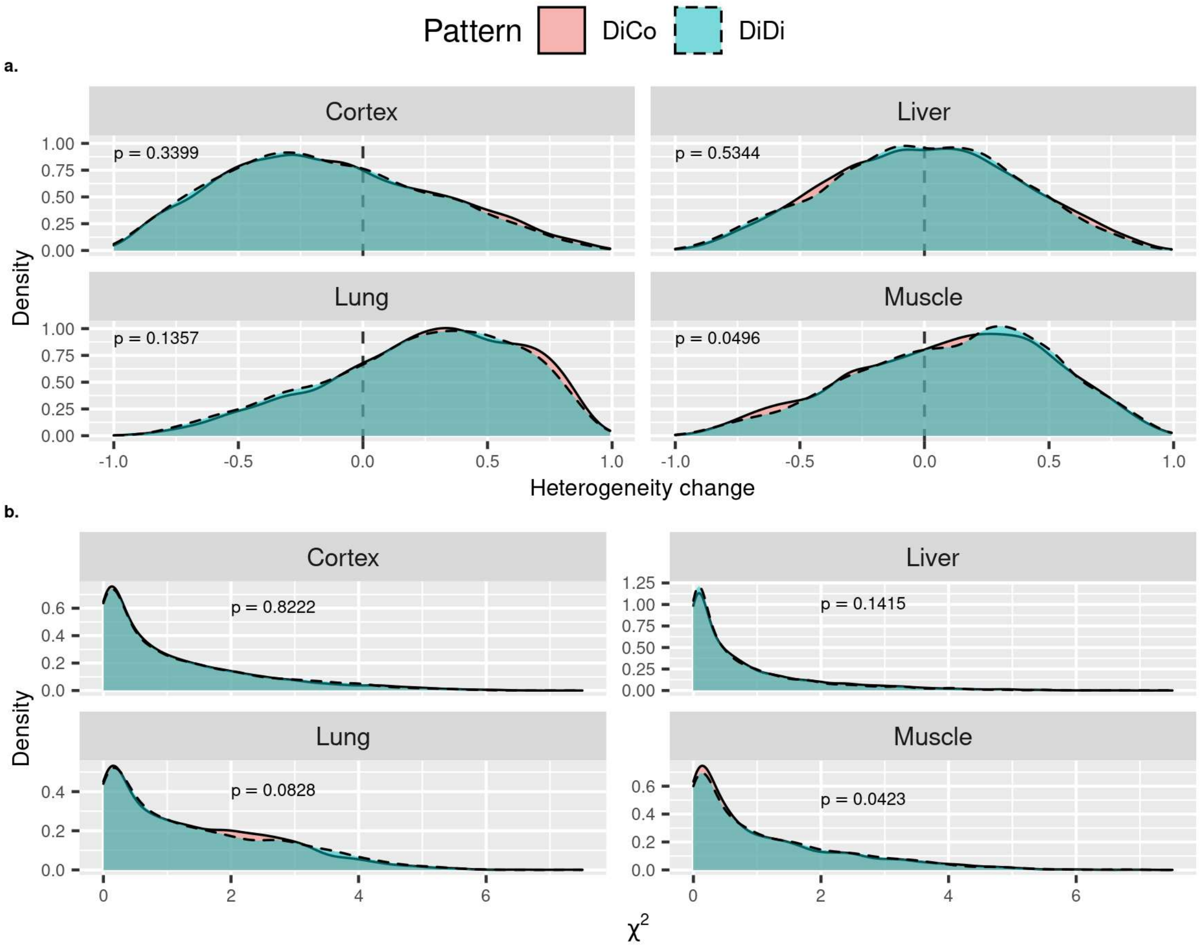


*Two different heteroscedasticity tests were performed to compare DiCo (n=4,802) vs DiDi (n=4,182, divergent throughout the lifetime) genes to test whether the convergence pattern is a result of the regression towards the mean.* ***a)*** *Density plots of Spearman's correlation coefficients (x-axis) between heterogeneity and age for DiCo and DiDi genes, in each tissue. Heterogeneity was calculated as the absolute residuals of the linear regression between age (log2 scale) and expression (see Methods). Only in muscle tissue, the two-sided Kolmogorov-Smirnov (KS) test result was marginally significant in the direction of higher heterogeneity change for DiDi genes (p = 0.0496).* ***b)*** *Density plots of Chi-Square test statistics (x-axis) from Breusch-Pagan test (from "car" package in R) between expression level and age (log2 scale) for DiCo and DiDi genes, in each tissue. Only in muscle tissue, the two-sided KS test result was significant in the direction of higher heterogeneity change for DiDi genes (p = 0.0423). P-values of KS test results between DiCo and DiDi genes are given within each plot.*

**Figure 2-figure supplement 16. Sex effect on CoV analysis using GTEx**

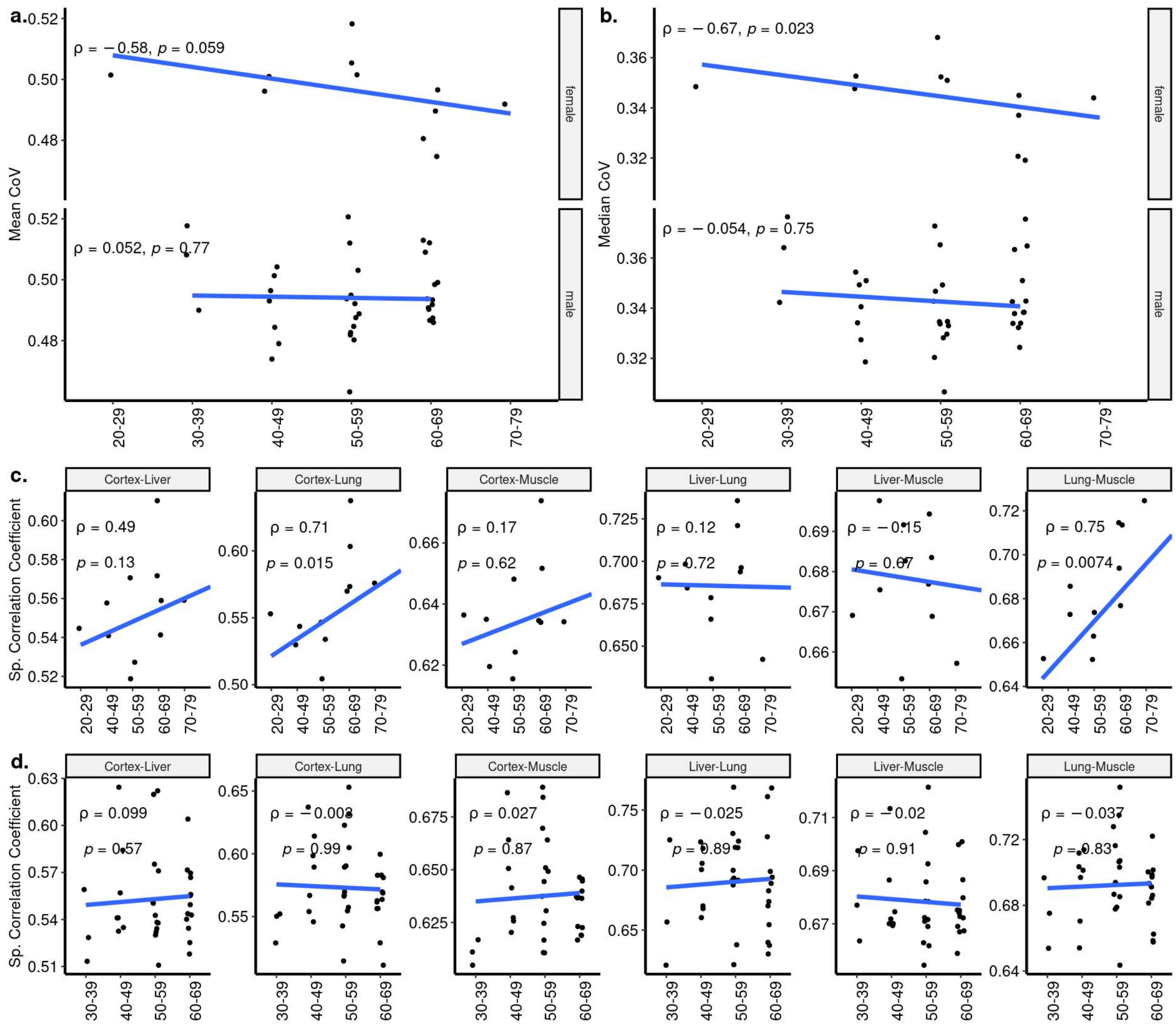


***a-b)** Transcriptome-wide mean **(a)** and median **(b)** CoV change with age across four tissues (Cortex, Liver, Lung, Muscle) in GTEx for female (n=11) and male (n=36) individuals, separately. Each point represents the mean or median CoV value of all protein-coding genes (16,197) for each individual. Spearman's correlation coefficients and p-values are also presented in the plots. **c-d)** The change in pairwise Spearman's correlation coefficient between gene expression values of the same individual (y-axis) for **(c)** females (n=11) and **(d)** males (n=36), across ages (x-axis). Spearman's correlation coefficient and p-values between the pairwise tissue correlations and age are also presented in each plot.*

**Figure 2-figure supplement 17. PCA of Schaum dataset covering cortex, liver, lung, and muscle tissues**

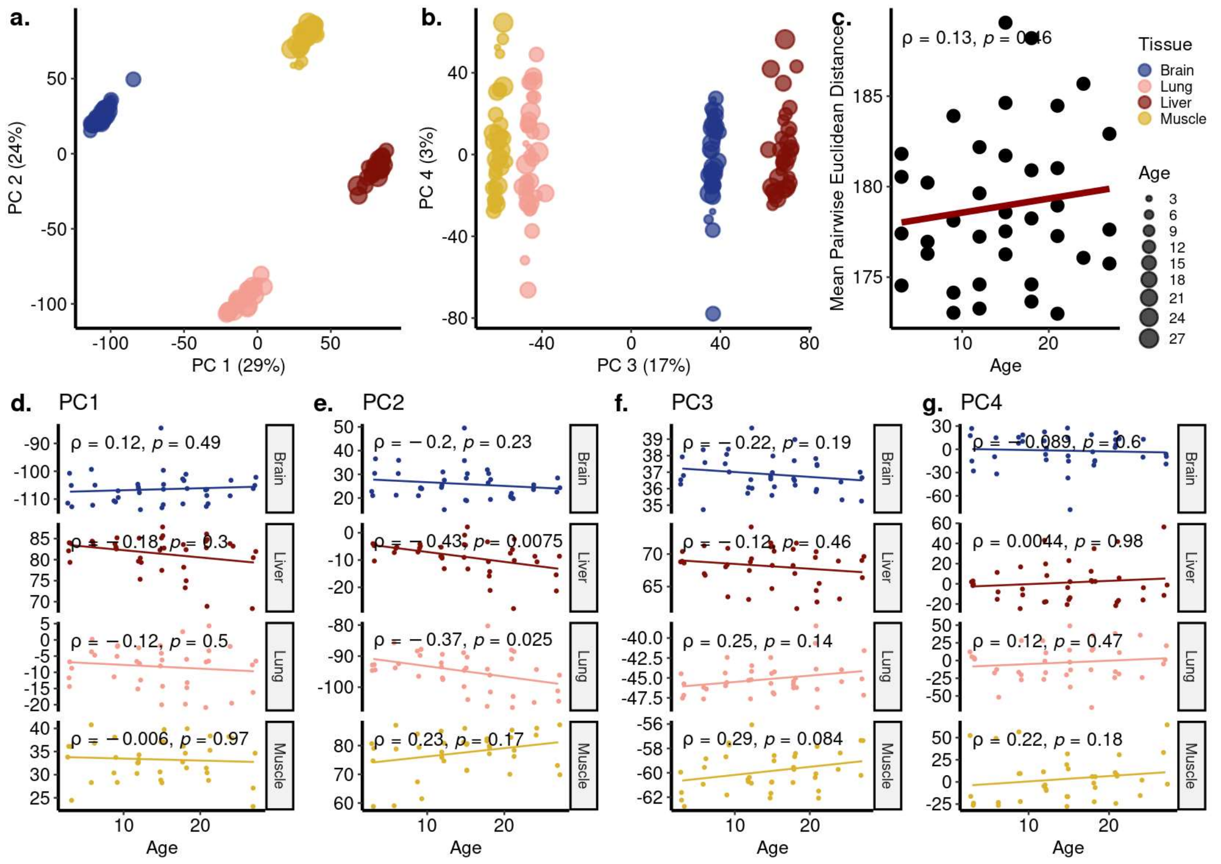


***a-b)*** *Principal components analysis (PCA) of expression values of 16,806 genes across four tissues (Cortex, Liver, Lung, Muscle) of 37 individuals in the Schaum dataset. Values in parentheses show the variance explained by each PC.* ***c)*** *The change in mean pairwise Euclidean distance between the PC values for the tissues of the same individuals (y-axis) with age (x-axis).* ***d-g)*** *Association between the first four PCs (y-axis) and age (x-axis). The tissue and age of the samples are indicated by the colour and size of the points, respectively. Spearman's correlation test results are indicated in each plot.*

**Figure 2-figure supplement 18. CoV and pairwise correlation analysis of Schaum dataset covering cortex, liver, lung, and muscle tissues**

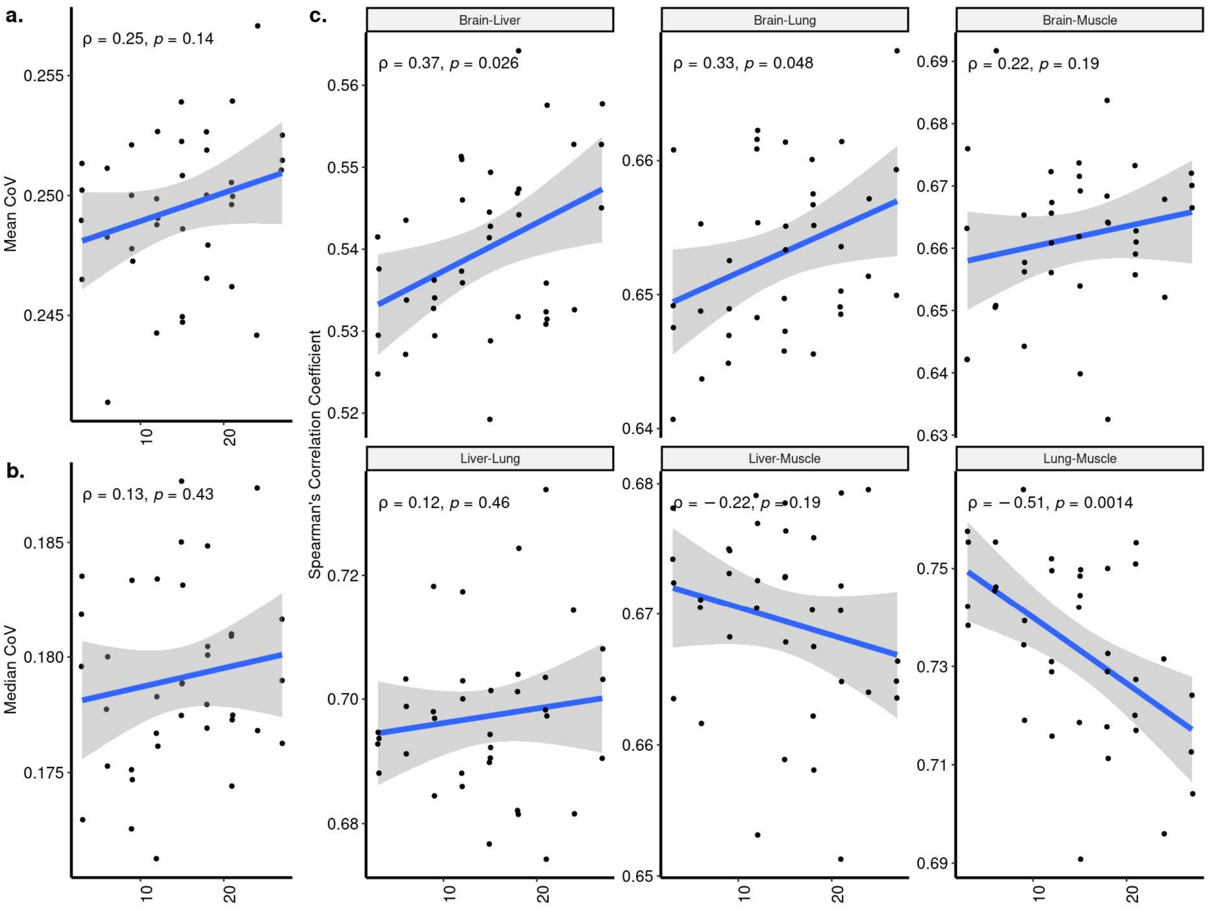


***a-b)*** *Transcriptome-wide mean* ***(a)*** *and median* ***(b)*** *CoV change with age across four tissues (Cortex, Liver, Lung, Muscle) in Schaum dataset. Each point represents the mean or median CoV value of all protein-coding genes (16,806) for each individual (n=37). Spearman's correlation coefficients and p-values are also presented in the plot.* ***c)*** *The change in pairwise Spearman's correlation coefficient between gene expression values of the same individual across ages (y-axis) with age (x-axis). Spearman's correlation coefficient and p-values between the pairwise tissue correlations and age are also presented in each plot.*

**Figure 2-figure supplement 19. PCA of Schaum dataset with eight tissues**

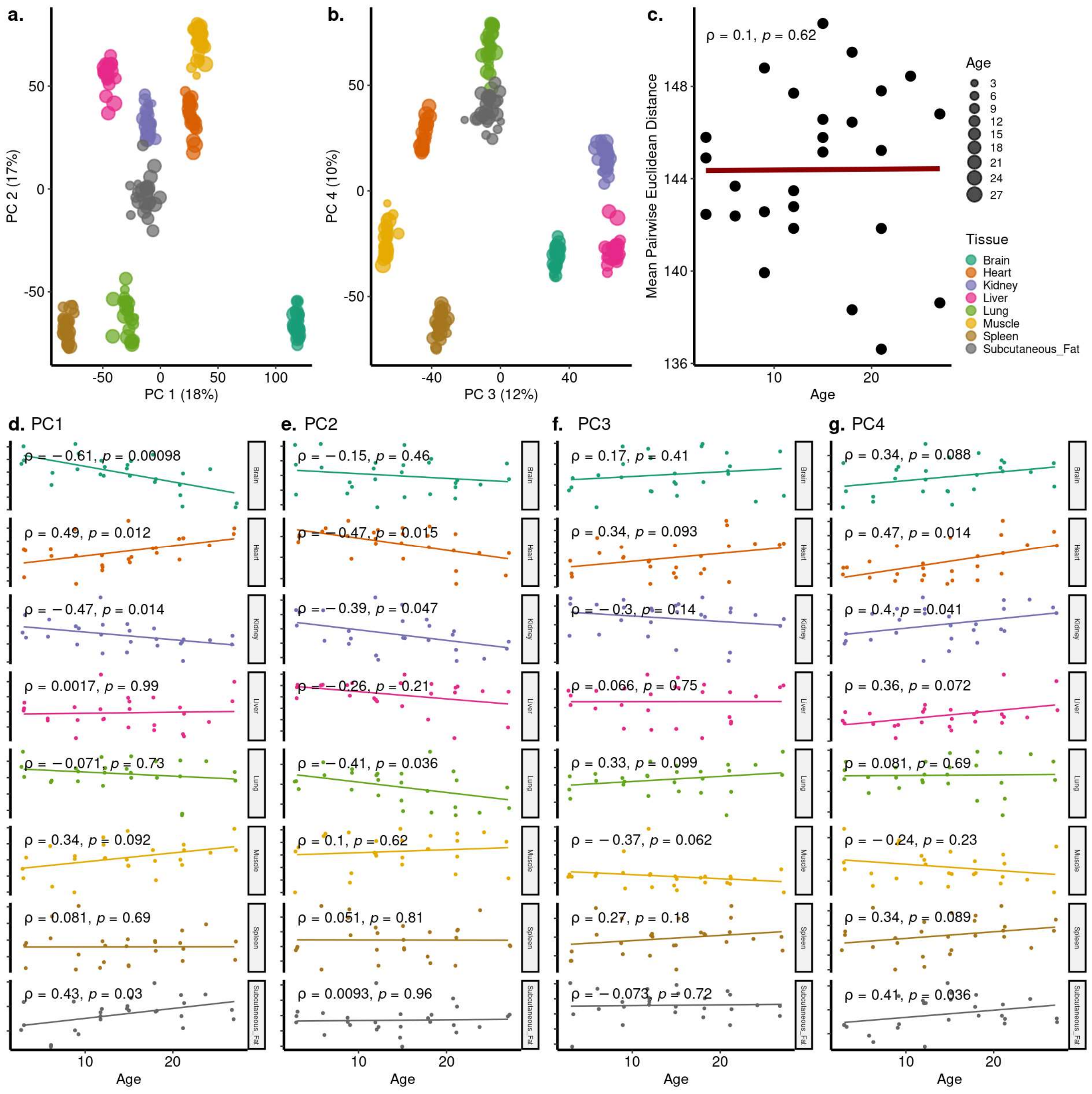


***a-b)** Principal components analysis (PCA) of expression values of 17,619 genes across eight tissues of 26 individuals in the Schaum dataset. Values in parentheses show the variance explained by each PC. **c)** The change in mean pairwise Euclidean distance between the PC values for the tissues of the same individuals (y-axis) with age (x-axis). **d-g)** Association between the first four PCs (y-axis) and age (x-axis). The tissue and age of the samples are indicated by the colour and size of the points, respectively.*

**Figure 2-figure supplement 20. CoV and pairwise correlation analysis of Schaum dataset with eight tissues**

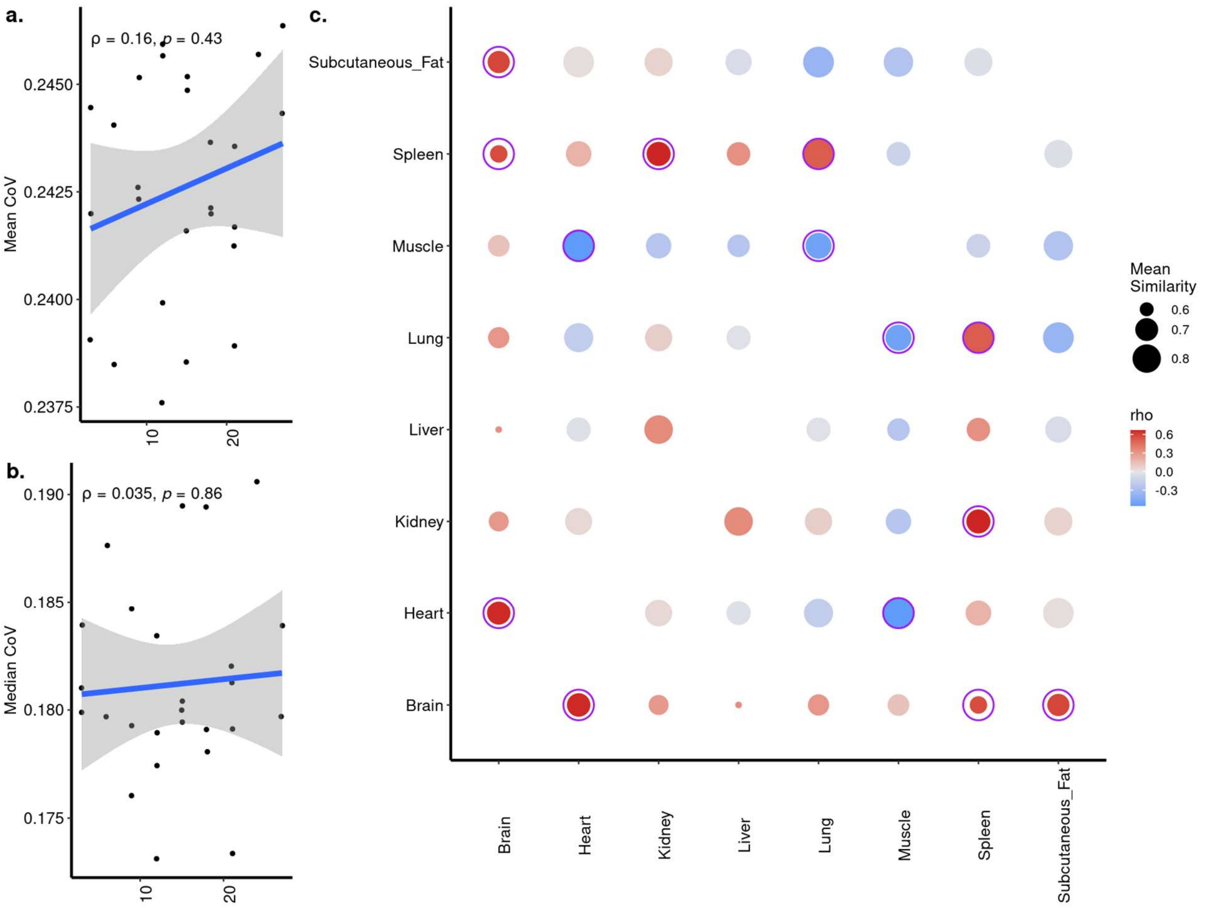


***a-b)*** *Transcriptome-wide mean **(a)** and median **(b)** CoV change with age across eight tissues (Brain (Cortex), Heart, Kidney, Liver, Lung, Muscle, Spleen, Subcutaneous Fat) in Schaum dataset. Each point represents the mean or median CoV value of all protein-coding genes (17,619) for each individual (n=26). Spearman's correlation coefficients and p-values are also presented in the plot.* ***c)*** *Age-related changes in pairwise Spearman's correlation coefficient between gene expression values of the same individual. The colour of points shows the correlations between age and pairwise correlations, where darker red colour indicates an increased correlation with age and darker blue indicates a decreased correlation. The size of points shows the mean similarity (correlation) between tissues using all ages. Significant correlations are indicated with circles around the points after multiple testing correction using 'BH'. (5/7 of significant correlations were positive).*

**Figure 4-figure supplement 1. Age-related expression change trends in DiCo enriched categories denoted as ‘Other GO’ in the first clustering**

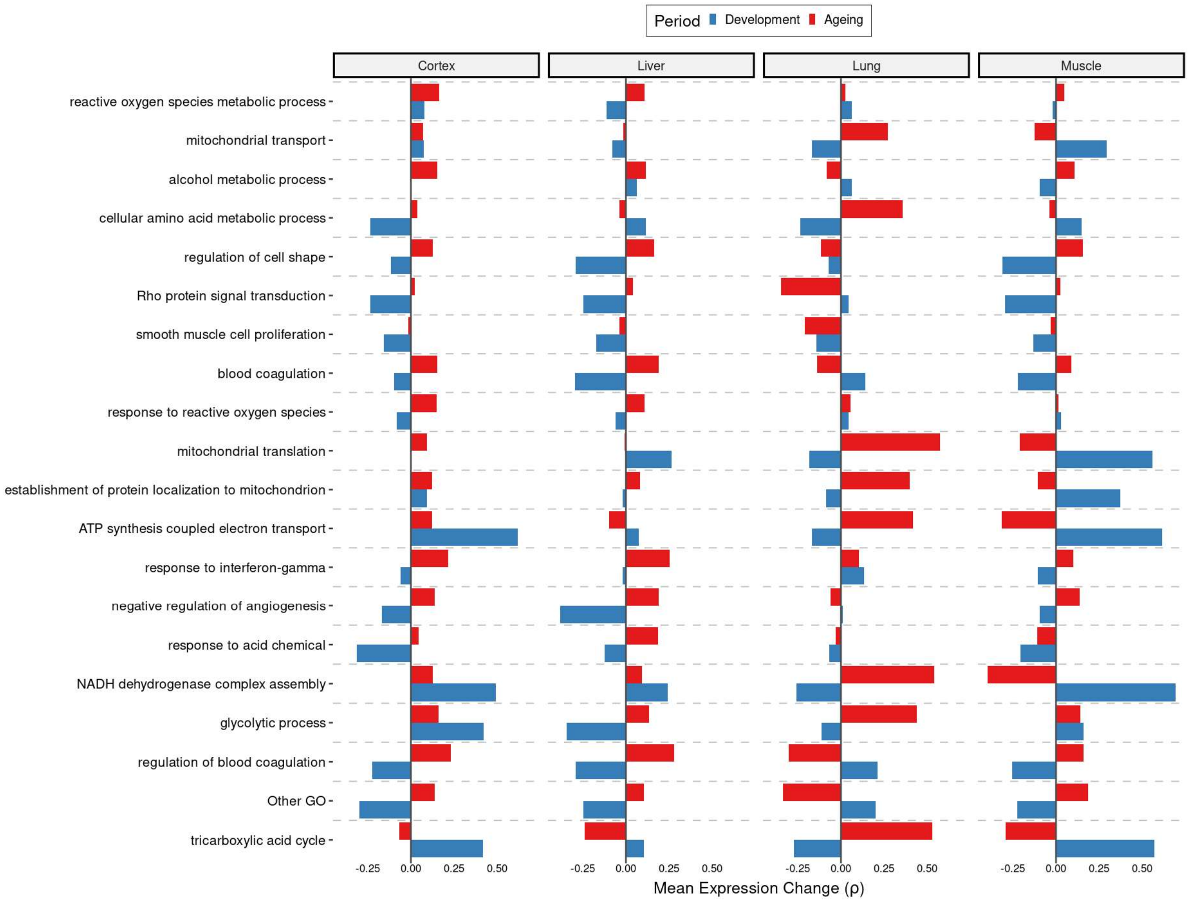


*Age-related expression change trends of genes (x-axis) in categories enriched in DiCo (GSEA) that were grouped into one cluster ‘Other GO’ in **Figure 4g**. These categories (n=69) were again summarised into representatives (y-axis) using hierarchical clustering and Jaccard similarities (see Methods). Categories are ordered by the number of genes they contain from highest (bottom, n = 97) to lowest (top, n = 21). One cluster containing unrelated categories (n=17) was again denoted as ‘Other GO’.*

**Figure 4-figure supplement 2. Comparison of datasets**

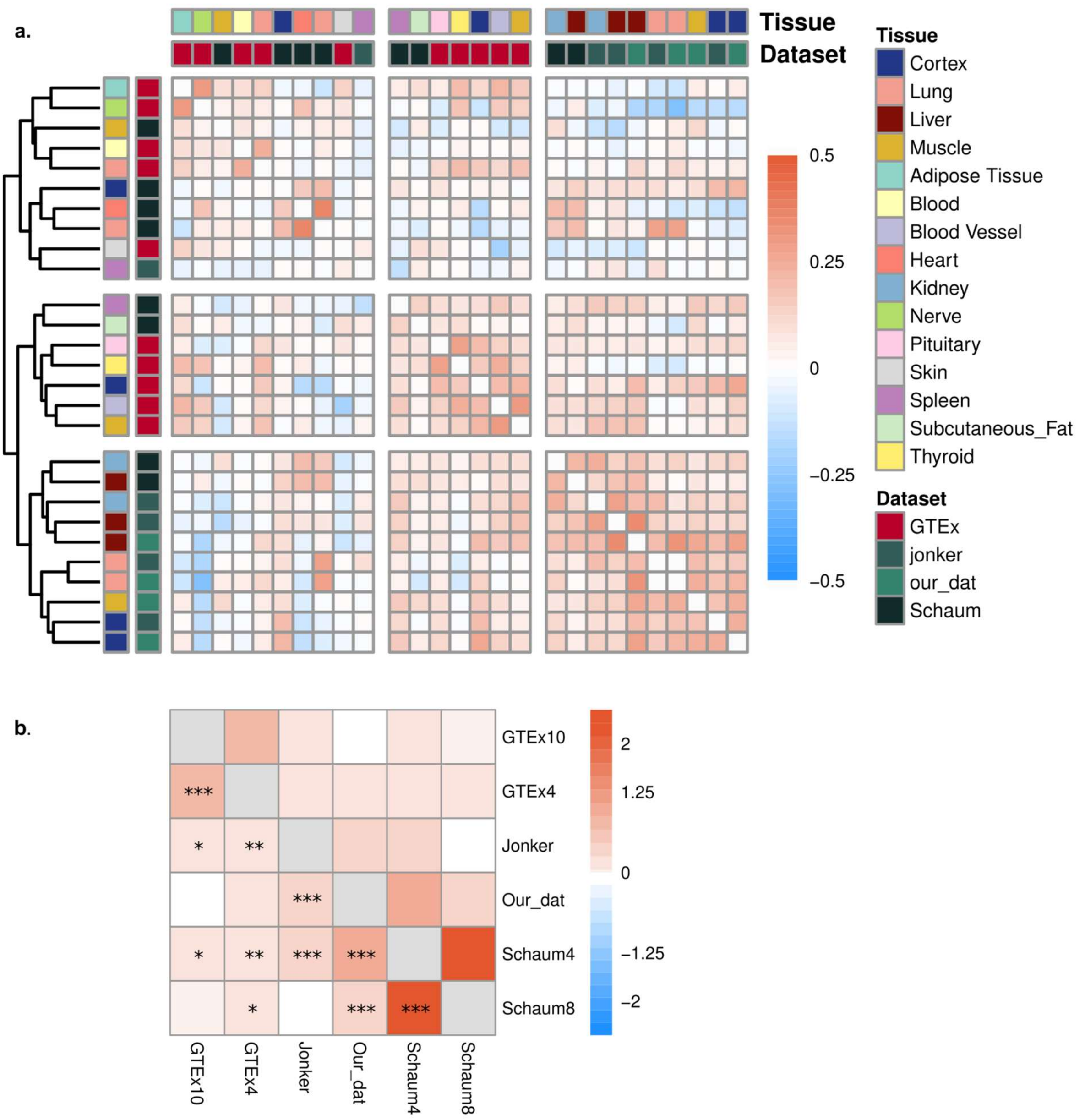


*a) Heatmap using Spearman's correlation coefficients among expression trajectories (Spearman's correlation coefficients between expression and age) across datasets during ageing. As the pairwise tissue correlations range between -0.2 to 0.52, the colour palette was restricted to -0.52 to 0.52 range. The same tissues of our dataset and Jonker dataset were clustered together (cortex, lung, liver) in the lower right corner.* ***b)*** *Enrichment of convergent genes among datasets during ageing. GTEx10 and GTEx4: CoV calculation was performed with ten tissues and with the same four tissues as our dataset in GTEx. Schaum8 and Schaum4: CoV calculation was performed with eight tissues and with the same four tissues as our dataset in Schaum dataset.'***': FDR corrected p-value<0.001, '**': FDR corrected p-*

*value<0.01, '*': FDR corrected p-value<0.1. All log2(OR) values were positive except for our data vs GTEx10 (log2(OR)= -0.04) and Jonker vs Schaum8 (log2(OR) = -0.06), both of which were non-significant.*

**Figure 5-figure supplement 1. Age-related changes in cell type proportions calculated using DiCo and non-DiCo genes**

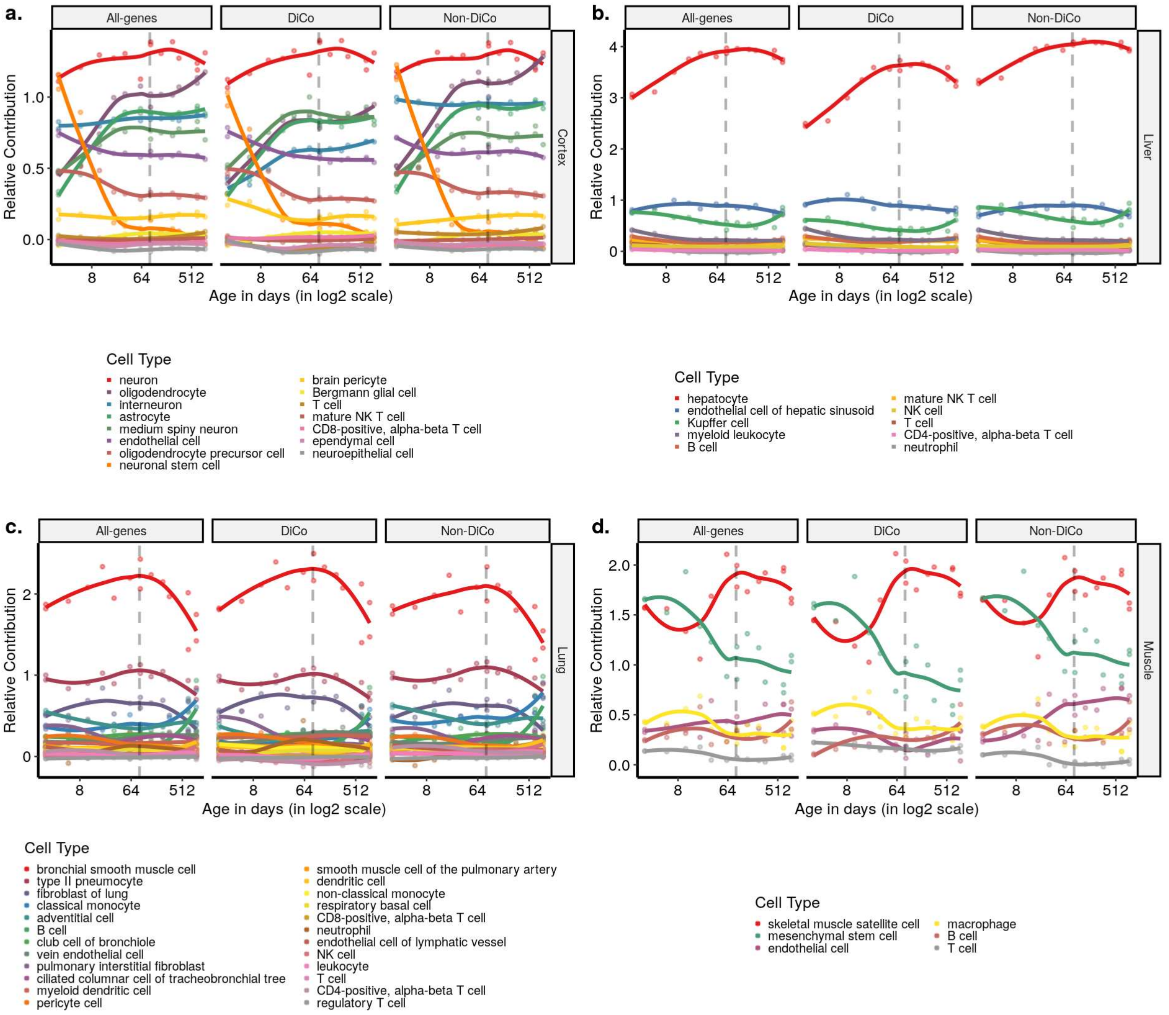


*Deconvolution of bulk tissue expression profiles of the mice in our dataset with regression analysis using the single-cell expression profile of the 3-month-old mice in the Tabula Muris Senis dataset. Contribution of each cell type was measured using three gene sets; all genes (n=[12,492, 12,849]), DiCo (n=[4,007, 4,106]) and non-DiCo genes (n=[8,485, 8,743]). Age-related changes of the relative contribution of each cell type in each tissue are given in* ***Figure 5-source data****.*

**Figure 5-figure supplement 2. Permutation-based comparison between DiCo and non-DiCo related cell type proportion changes with age in the cortex**

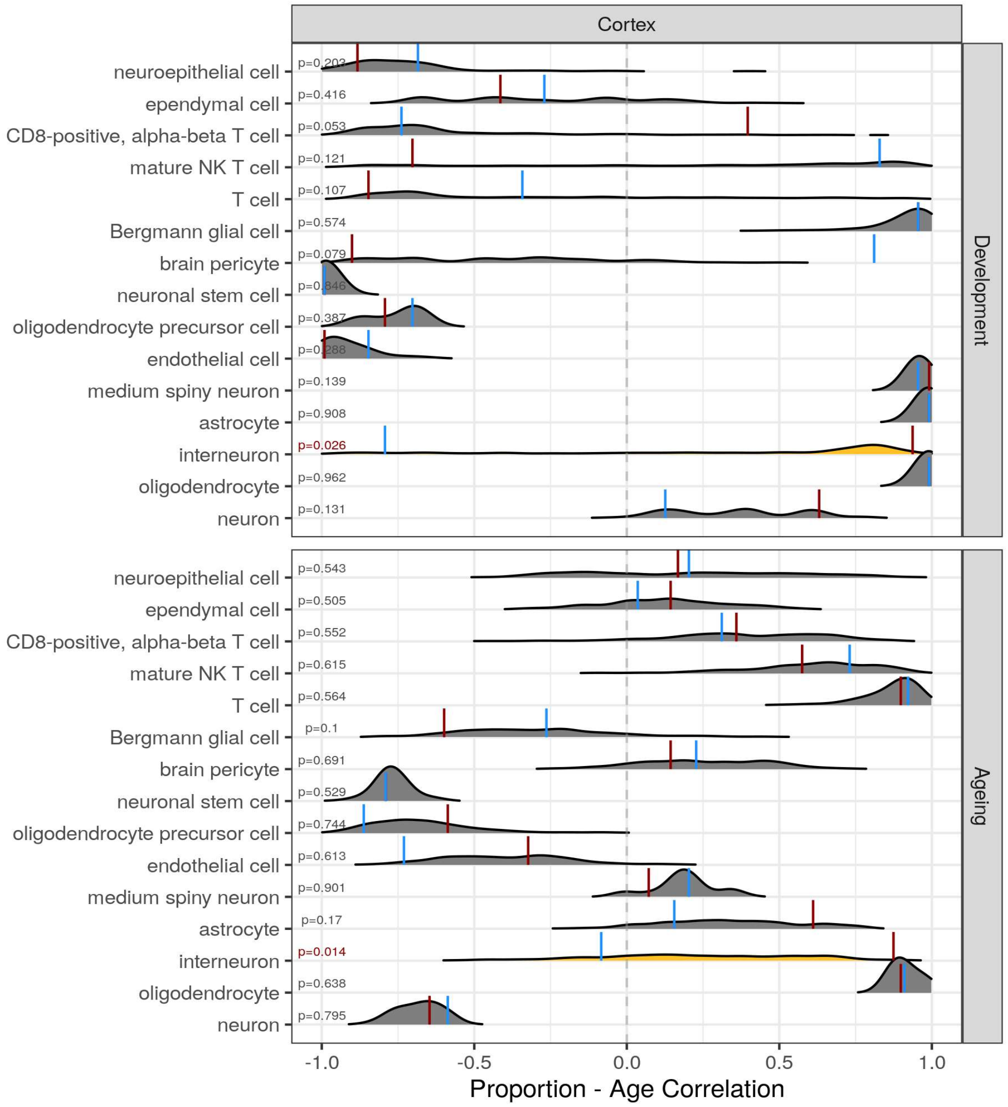


*The difference between DiCo (4,106) and non-DiCo (8,743) related cell type proportion changes with age was tested in the cortex tissue. The x-axis is the Spearman's correlation coefficient between age and relative contribution of a given cell type. The red vertical lines show the cell type proportion changes calculated with DiCo genes (observed value) and the blue vertical lines indicate the same but with non-DiCo genes. Overlapping DiCo and non-DiCo values were indicated with blue. Null distributions for non-DiCo genes (density plots) were created with re-sampling among all genes (n=12,849) (Methods). Significant results were represented with yellow density plots and the nominal p-values for permutation tests are indicated on the left side of the density plots. Permutation test results are also provided in **Figure 5-source data**.*

**Figure 5-figure supplement 3. Permutation-based comparison between DiCo and non-DiCo related cell type proportion changes with age in the liver**

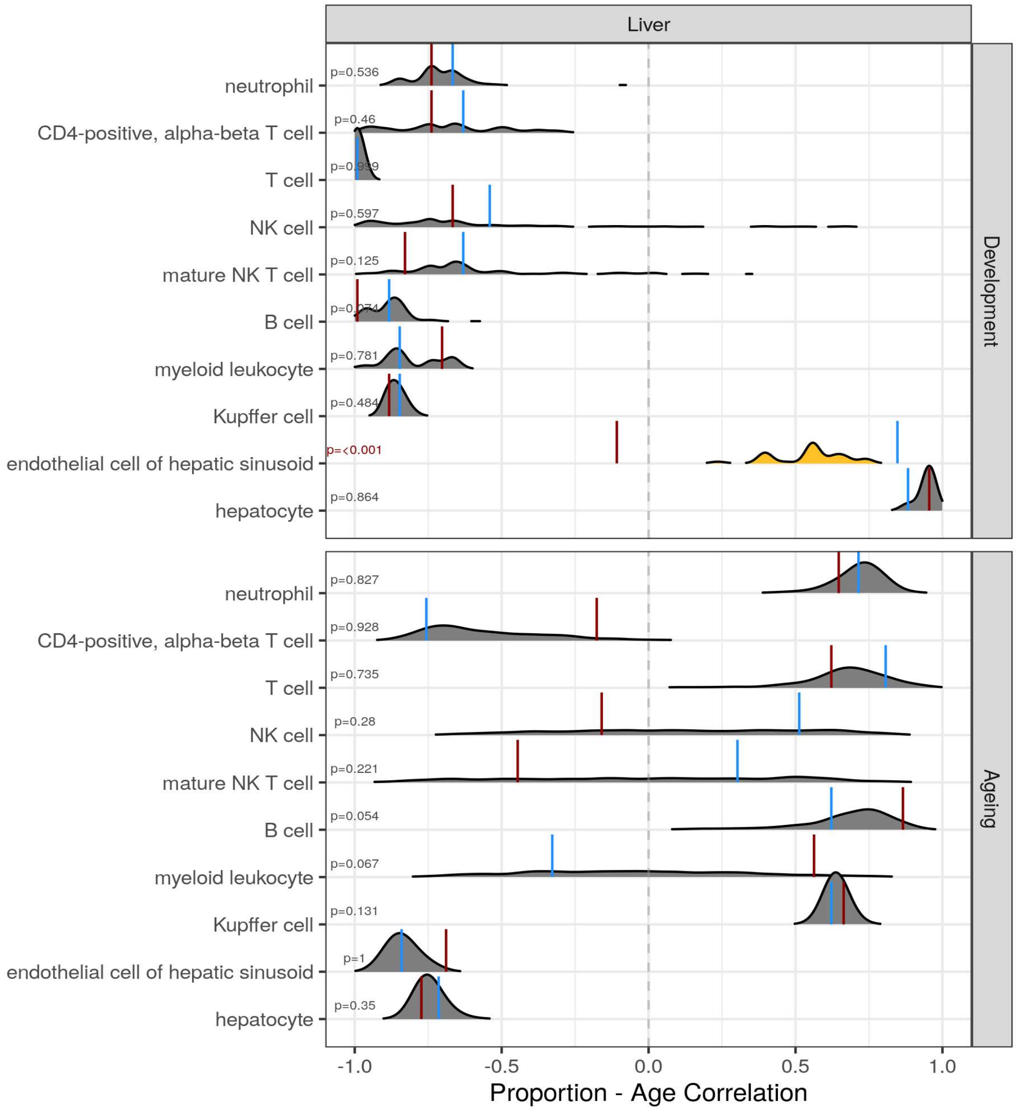


*The difference between DiCo (4,007) and non-DiCo (8,485) related cell type proportion changes with age was tested in the liver tissue. The x-axis is the Spearman's correlation coefficient between age and relative contribution of a given cell type. The red vertical lines show the cell type proportion changes calculated with DiCo genes and the blue vertical lines indicate the same but with non-DiCo genes. Overlapping DiCo and non-DiCo values were indicated with blue. Null distributions for non-DiCo genes (density plots) were created with re-sampling among all genes (n=12,492) (see Methods). Significant results were represented with yellow density plots and the nominal p-values for permutation tests are indicated on the left side of the density plots. Permutation test results are provided in* ***Figure 5-source data****.*

**Figure 5-figure supplement 4. Permutation-based comparison between DiCo and non-DiCo related cell type proportion changes with age in the lung**

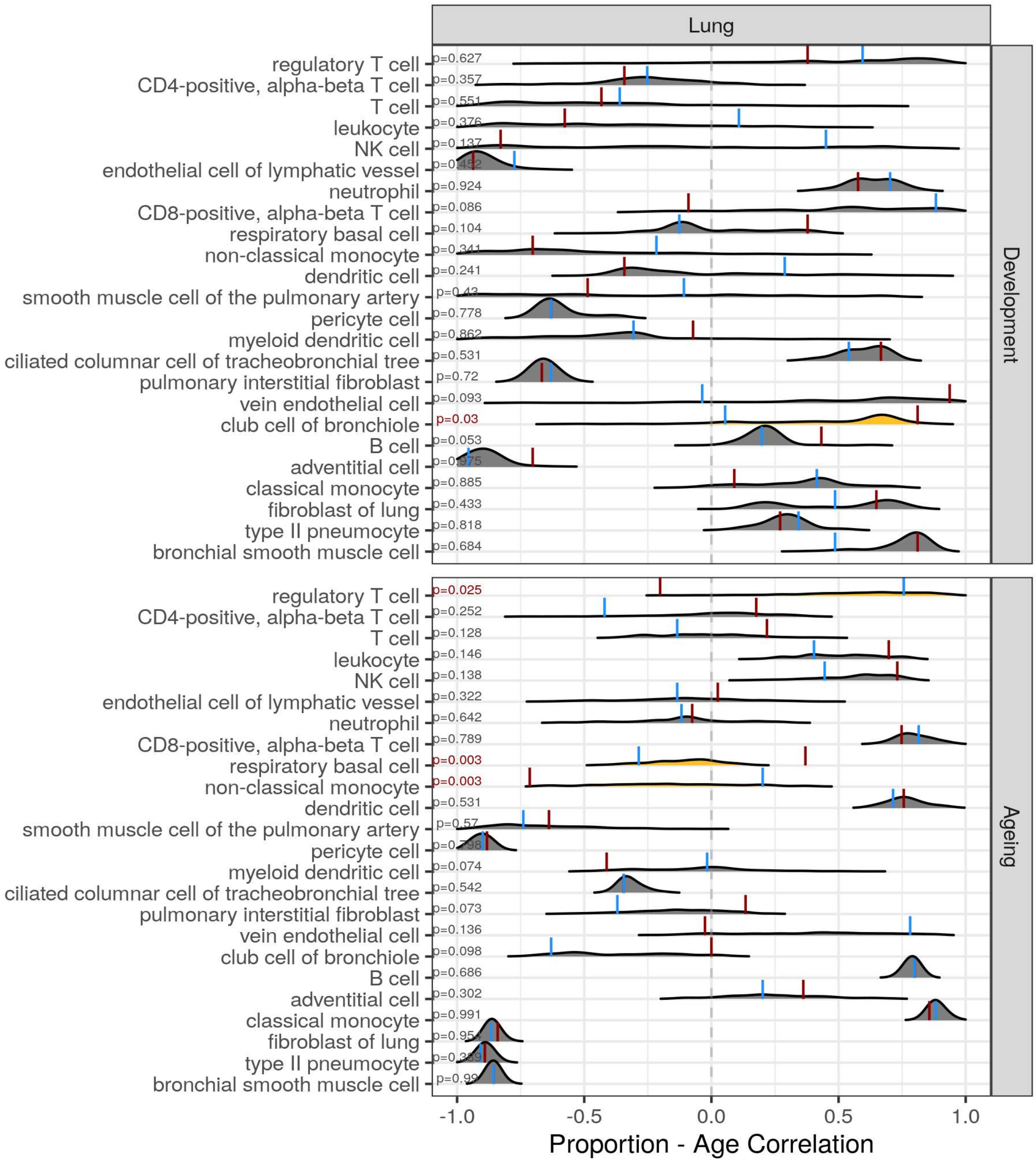


*The difference between DiCo (4,084) and non-DiCo (8,670) related cell type proportion changes with age was tested in the lung tissue. The x-axis is the Spearman's correlation coefficient between age and relative contribution of a given cell type. The red vertical lines show the cell type proportion changes calculated with DiCo genes and the blue vertical lines indicate the same but with non-DiCo genes. Overlapping DiCo and non-DiCo values were indicated with blue. Null distributions for non-DiCo genes (density plots) were created with re-sampling among all genes (n=12,754) (see*

*Methods). Significant results were represented with yellow density plots and the nominal p-values for permutation tests are indicated on the left side of the density plots. Permutation test results are provided in* ***Figure 5-source data****.*

**Figure 5-figure supplement 5. Permutation-based comparison between DiCo and non-DiCo related cell type proportion changes with age in the muscle.**

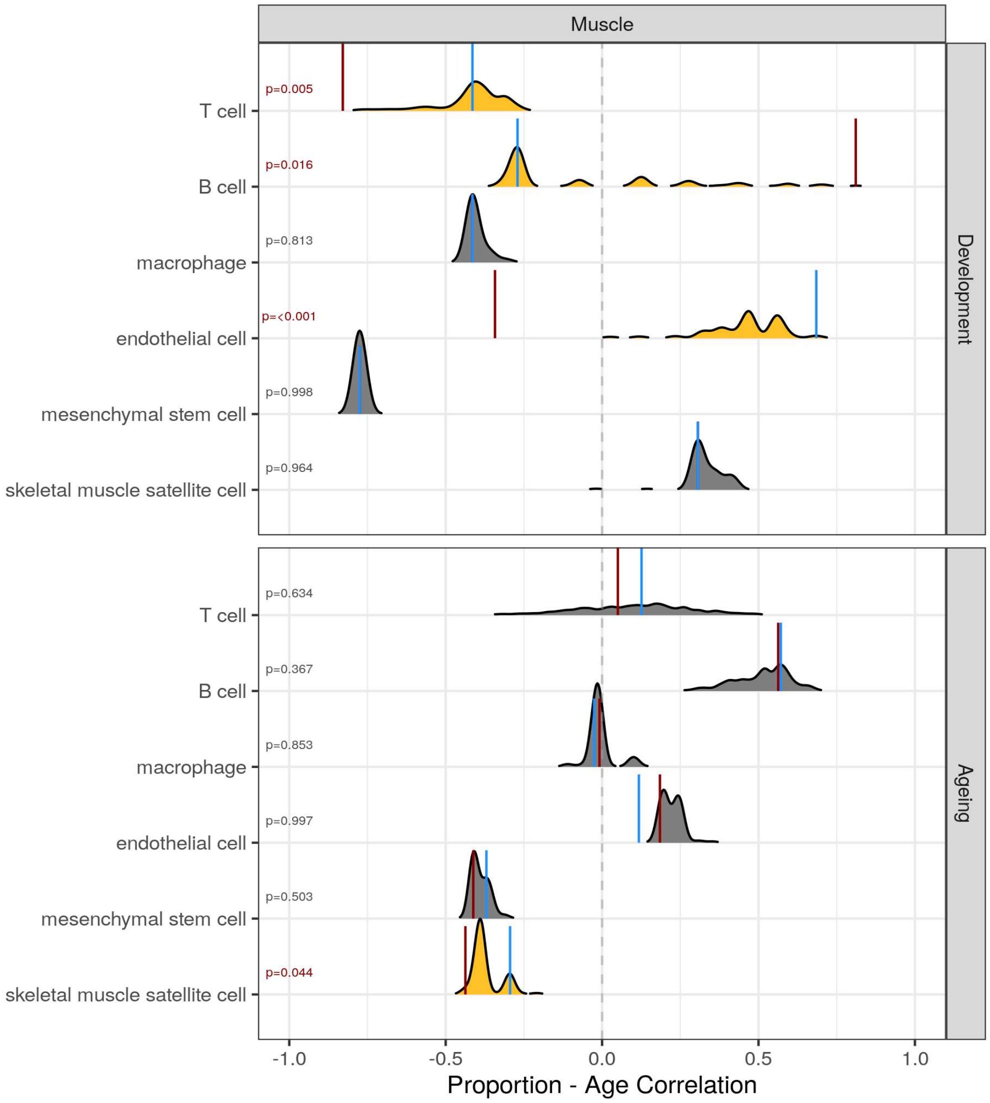


*The difference between DiCo (4,055) and non-DiCo (8,568) related cell type proportion changes with age was tested in the muscle tissue. The x-axis is the Spearman's correlation coefficient between age and relative contribution of a given cell type. The red vertical lines show the cell type proportion changes calculated with DiCo genes and the blue vertical lines indicate the same but with non-DiCo genes. Overlapping DiCo and non-DiCo values were indicated with blue. Null distributions for non-DiCo genes (density plots) were created with re-sampling among all genes (n=12,623) (see Methods). Significant results were represented with yellow density plots and the nominal p-values for permutation tests are indicated on the left side of the density plots. Permutation test results are provided in* ***Figure 5-source data****.*

**Figure 5-figure supplement 6. Intra-tissue CoV changes between cell types using Tabula Muris Senis dataset**

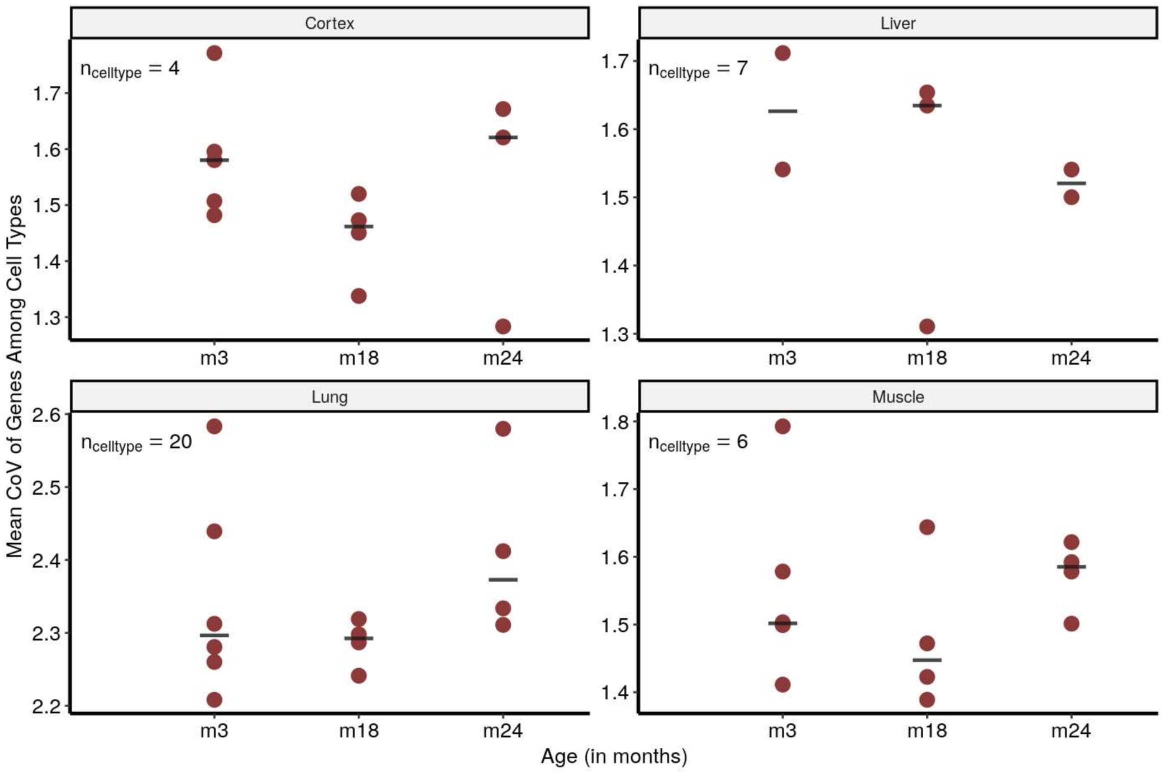


*Intra-tissue CoV: CoV is calculated among cell types within each tissue for each individual mouse and in 3 age groups. Y-axis shows the mean CoV value of genes for each individual. The horizontal line on each age group shows the median of points. Cell types found in at least 2 individuals at every time point were considered.*

### Source Data Files

**Figure 1-source data. Data summary, age-related expression patterns and reversal patterns.**

**Figure 2-source data. All the data related to DiCo pattern: age-related CoV change of genes, pairwise tissue expression correlations, analysis of independent datasets; GSE34378 (Jonker et al.), GSE132040 (Schaum et al.) and GTEx.**

**Figure 3-source data. Effect sizes for determination of tissue-specific genes, enrichment of DiCo and reversal genes within tissue-specific genes.**

**Figure 4-source data. GSEA result of DiCo genes, DiCo enrichment with tissue specific expression loss, age-related expression change correlations and convergence overlaps among datasets**

**Figure 5-source data. Cell type proportion estimation and cell-autonomous changes using Tabula Muris Senis dataset**

### Supplementary Files

#### Supplementary File 1. GORA of age-related genes in tissues

Tissue-specific age-related gene expression changes and functional enrichment test results, performed with gene over-representation analysis (GORA) using 'topGO' package.

#### Supplementary File 2. GORA of shared age-related genes among tissues

Functional enrichment for shared genes across tissues. The same GORA that was performed for Supplementary File 1, was used to test the enrichment of shared up/down-regulated genes in development among the background genes which are chosen as the all significant age-related genes across tissues in development. We did not apply the test for the ageing period as there were no shared ageing-related expression changes.

#### Supplementary File 3. GORA of reversal patterns

Functional enrichment for gene expression reversals. GORA analysis was performed with the same criteria as explained above. Up-Down reversal genes were tested against Up-Up genes and Down-Up reversal genes were tested against Down-Down genes in each tissue.

#### Supplementary File 4. GORA of DiCo gene clusters determined with CoV values

Functional enrichment of DiCo genes clustered with kmeans algorithm according to their CoV values. GORA analysis was performed using gene sets in each cluster (Figure 2–figure supplement 2) which were tested among all DiCo genes.

#### Supplementary File 5. GORA of DiCo gene clusters determined with expression levels

Functional enrichment of DiCo genes clustered with kmeans algorithm according to their expression levels. Gora analysis was performed using gene sets in each cluster (Figure 2–figure supplement 3) which are tested among all DiCo genes.
